# Systems analysis reveals ageing-related perturbations in retinoids and sex hormones in Alzheimer’s and Parkinson’s diseases

**DOI:** 10.1101/2021.06.10.447367

**Authors:** Simon Lam, Nils Hartmann, Rui Benfeitas, Cheng Zhang, Muhammad Arif, Hasan Turkez, Mathias Uhlén, Christoph Englert, Robert Knight, Adil Mardinoglu

**Affiliations:** Faculty of Dentistry, Oral and Craniofacial Sciences, King’s College London, London, SE1 9RT, United Kingdom; Leibniz Institute on Aging, Fritz Lipmann Institute, Jena, 07745, Germany; National Bioinformatics Infrastructure Sweden (NBIS), Science for Life Laboratory, Department of Biochemistry and Biophysics, Stockholm University, Stockholm, SE-17121, Sweden; Science for Life Laboratory, KTH – Royal Institute of Technology, Stockholm, SE-17121, Sweden; Department of Medical Biology, Faculty of Medicine, Atatürk University, Erzurum, 25240, Turkey; Institute of Biochemistry and Biophysics, Freidrich-Schiller-University Jena, Jena, 07745, Germany

## Abstract

Neurodegenerative diseases (NDDs), including Alzheimer’s (AD) and Parkinson’s diseases (PD), are complex heterogeneous diseases with highly variable patient responses to treatment. Due to the growing evidence for ageing-related clinical and pathological commonalities between AD and PD, these diseases have recently been studied in tandem. In this study, we analyse transcriptomic data from AD and PD patients, and stratify these patients into three subclasses with distinct gene expression and metabolic profiles. Through integrating transcriptomic data with a genome-scale metabolic model and validating our findings by network exploration and co-analysis using a zebrafish ageing model, we identify retinoids as a key ageing-related feature in all subclasses of AD and PD. We also demonstrate that the dysregulation of androgen metabolism by three different independent mechanisms is a source of heterogeneity in AD and PD. Taken together, our work highlights the need for stratification of AD/PD patients and development of personalised and precision medicine approaches based on the detailed characterisation of these subclasses.

## Introduction

Neurodegenerative diseases (NDDs), including Alzheimer’s (AD) and Parkinson’s diseases (PD), cause years of a healthy life to be lost. Much previous AD and PD research has focused on the causative neurotoxicity agents, namely amyloid β and α-synuclein, respectively. The current front-line therapies for AD and PD are cholinesterase inhibition and dopamine repletion, respectively, which are considered gold standards. Unfortunately, these therapies are not capable of reversing neurodegeneration (Liberini et al., 1996; Wijemanne and Jankovic, 2015), thus necessitating potentially lifelong dependence on the drug and risking drug-associated complications. Moreover, AD and PD are complex diseases with heterogeneous underlying molecular mechanisms involved in their progression (Greenland et al., 2019; Long and Holtzman, 2019). This variability can explain the differences in patient response to other treatments such as oestrogen replacement therapy (Baum, 2005; Meoni et al., 2020) and statin treatment (Shepardson et al., 2011; Jeong et al., 2019). Hence, we observed that there are distinct disease classes affecting specific cellular processes. Therefore, there is a need for the development of personalised treatment regimens.

In this study, we propose a holistic view of the mechanisms underlying the development of NDDs rather than focusing on amyloid β and α-synuclein (Lam et al., 2020). To date, complex diseases including liver disorders and certain cancers have been well studied through the use of metabolic modelling. This enabled the integration of multiple omics data for stratification of patients, discovery of diagnostic markers, identification of drug targets, and proposing of personalised or class-specific treatment strategies (Mardinoglu et al., 2018; Altay et al., 2019; Joshi et al., 2020; Lam et al., 2021). A similar approach may be applied for AD and PD since there is already a wealth of data from AD and PD patients from postmortem brain tissues and blood transcriptomics.

AD and PD share multiple clinical and pathological similarities, including comorbidities (Stampfer, 2006; De La Monte and Wands, 2008), inverse associations with cancer (Bajaj et al., 2010; Driver et al., 2012), and ageing as a risk factor (Hindle, 2010; Sengoku, 2020). One type of ageing is telomeric ageing, which is associated with the loss of telomeres, protein/nucleic acid structures that protect chromosome ends from degradation (Chakravarti et al., 2021). The enzyme telomerase is necessary for the maintenance of telomeres. In adults, telomerase activity is mostly limited to progenitor tissues such as in the ovaries, testes, and bone marrow. Loss of telomerase activity leads to telomere shortening, loss of sequences due to end-replication, and eventual degradation of sequences within coding regions, leading to telomeric ageing. Considering NDDs as a product of ageing, we can use an ageing model organism to study its effects on the brain. In our study, we used zebrafish (*Danio rerio*) as model organism since it has been used extensively used to study vertebrate ageing (Carneiro et al., 2016). For example, a zebrafish ageing model can harbour a nonsense mutation in the *tert* gene, which encodes the catalytic subunit of telomerase, and exhibit faster-than-normal ageing (Anchelin et al., 2013; Henriques et al., 2013).

In our study, we first analysed postmortem brain gene expression data and protein-protein interaction data from the Genotype-Tissue Expression (GTEx) database (GTEx Consortium, 2013), Functional Annotation of the Mammalian Genome 5 (FANTOM5) database (Forrest et al., 2014; Lizio et al., 2015, 2019; Marbach et al., 2016), Human Reference Protein Interactome (HuRI) database (Luck et al., 2019) and Human Protein Atlas (HPA) [http://www.proteinatlas.org, accessed 2021-03-09] (Uhlén et al., 2015) for characterization of normal brain tissue (**Figure 1A**). Secondly, we analysed transcriptomic data from the Religious Orders Study and Rush Memory Aging Project (ROSMAP) (Myers et al., 2007; Webster et al., 2009; Mostafavi et al., 2018) with published expression data from anterior cingulate cortices and dorsolateral prefrontal cortices of PD and Lewy body dementia patients, hereafter referred to as the Rajkumar dataset (Rajkumar et al., 2020), and from putamina, substantiae nigrae, and prefrontal cortices from patients with PD, hereafter referred to as the Zhang/Zheng dataset (Zhang et al., 2005; Zheng et al., 2010). On these data, we conducted differential gene expression and functional analysis, and then constructed biological networks to further explore coordinated patterns of gene expression. Next, we performed global metabolic analyses using genome-scale metabolic modelling. Alongside these analyses, we also leveraged zebrafish *tert* mutants to test the hypothesis that the identified changes may be associated with an ageing mechanism. Finally, based on our integrative systems analysis, we define three distinct disease subclasses within AD and PD and identified retinoids as a common feature of all three subclasses and likely to be perturbed through ageing. We reveal subclass-specific perturbations at three separate processes in the androgen biosynthesis and metabolism pathway, namely oestradiol metabolism, cholesterol biosynthesis, and testosterone metabolism.

**Figure 1.**
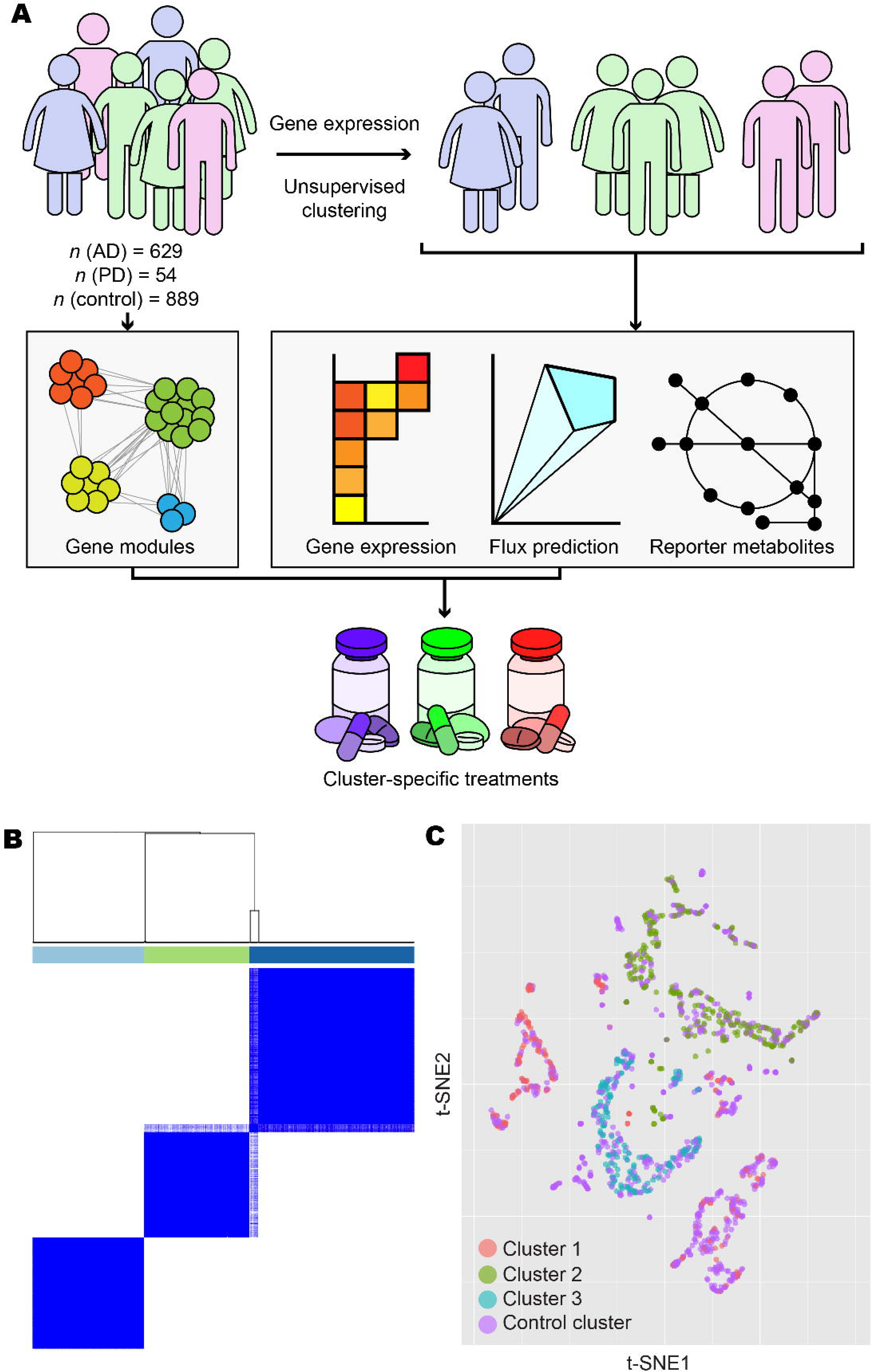
Overview and exploratory data analysis. **A)** Workflow for the analysis of human AD and PD samples. **B)** AD and PD samples were clustered into *k* clusters without supervision on the basis of normalised expression counts. Results are shown *k* = 3 and 1000 bootstrap replicates. Colour bars indicate cluster identity for each sample. For 2 ≤ *k* ≤ 7, refer to Supplementary figure 1. **C**) Normalised expression data from AD, PD, and control samples were projected onto 2-D space using t-distributed stochastic neighbour embedding (t-SNE). Points are coloured according to cluster assignment by unsupervised clustering. For further data visualisation, refer to Supplementary figure 2.

## Results

### Stratification of patients reveals three distinct disease classes

We retrieved gene expression and protein-protein interaction data from GTEx, FANTOM5, HuRI, HPA, and ROSMAP databases and integrated these data with the published datasets by Rajkumar and Zhang/Zheng. After performing quality control and normalisation (Materials and Methods), a total of 629 AD samples, 54 PD samples, and 889 control samples were included in the analysis (**Table 1**). To reveal transcriptomic differences between AD/PD samples compared to healthy controls, we identified differentially expressed genes (DEGs) and performed gene set enrichment (GSE) analyses. However, since AD and PD are complex diseases with no single cure, it is likely that multiple gene expression profiling exist, manifesting in numerous disease classes requiring distinct treatment strategies. We therefore used unsupervised clustering to elucidate these expression profiles and stratify the AD and PD patients based on the underlying molecular mechanisms involved in the disease occurrence.

**Table 1.** Summary of expression data sources. Expression data from AD and PD samples were obtained from the Genotype-Tissue Expression (GTEx) database, Functional Annotation of the Mammalian Genome 5 (FANTOM5) database, Human Protein Atlas (HPA), Religious Orders Study and Rush Memory Aging Project (ROSMAP), Rajkumar dataset, and Zhang/Zheng dataset.

| **Source** | **AD samples** | **PD samples** | **Control samples** |
| --- | --- | --- | --- |
| GTE<sub>x</sub>/FANTOM5 | 0 | 0 | 67 |
| HPA | 0 | 0 | 52 |
| Rajkumar | 0 | 14 | 13 |
| ROSMAP | 629 | 0 | 704 |
| Zhang/Zheng | 0 | 40 | 53 |
| Total | 629 | 54 | 889 |

Following unsupervised clustering with ConsensusClusterPlus (Wilkerson and Hayes, 2010), AD and PD samples separated into three clusters (**Figure 1B**, **Supplementary figure 1**). Clusters 1 and 2 contained samples from Zhang/Zheng and Rajkumar datasets, respectively, in addition to samples in the ROSMAP dataset. Cluster 3 contained only ROSMAP samples. Clusters did not form firmly along lines of sex, age, or brain tissues or brain subregion (**Supplementary figure 2**). Samples from non-diseased individuals were artificially added as a fourth, control cluster.

By differential expression analysis using DESeq2 (Love et al., 2014), we then characterised the distinct transcriptomic profiles within our disease clusters (**Figure 2A**). Cluster 1 showed mixed up- and downregulation of genes compared to control, whereas cluster 2 showed more downregulation and cluster 3 showed vast downregulation of genes compared to control.

**Figure 2.**
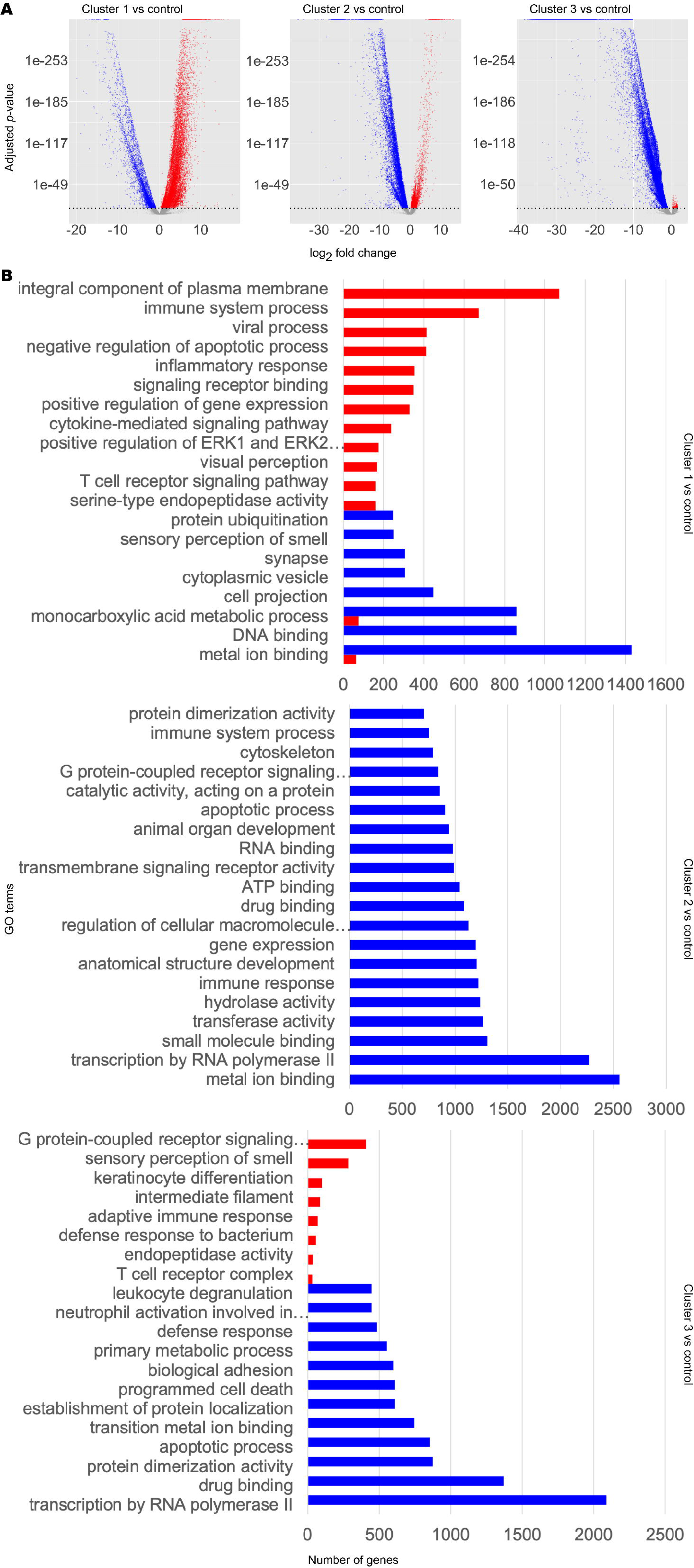
Transcriptomic and functional characterisation of AD and PD subclasses. Differentially expressed gene (DEG) analysis and gene set enrichment (GSE) analysis were performed for AD and PD and control samples for each disease cluster, using the control cluster as reference. **A)** DEG results. Significant DEGs were determined as those with a Benjamini-Hochberg adjusted *p*-value at or below a cut-off of 1×10^-10^. Upregulated significant DEGs are coloured red. Downregulated significant DEGs are coloured blue. Non-significant DEGs are coloured grey. **B)** Selected significantly enriched GO terms by number of genes as determined by GSE analysis. Red bars indicate upregulated GO terms. Blue bars indicate downregulated GO terms. For full data, refer to Supplementary data 1.

To infer the functional differences between the subclasses, we performed GSE analysis using piano (Väremo et al., 2013) (**Figure 2B**, **Supplementary data 1**). Globally, DEGs in any cluster 1-3 were enriched in upregulated Gene Ontology (GO) terms for immune response, olfaction, retinoid function, and apoptosis, but downregulated for copper ion transport and telomere organisation, compared to the control cluster. Considering individual clusters, cluster 1 DEGs were enriched in upregulated GO terms associated with immune signalling, cell signalling, and visual perception. We also found downregulation of GO terms associated with olfactory signalling and cytoskeleton. DEGs in cluster 2 were found to be enriched in downregulated GO terms associated with the cytoskeleton, organ development, cell differentiation, retinoid metabolism and response, DNA damage repair, inflammatory response, telomere maintenance, unfolded protein response, and acetylcholine biosynthesis and binding. On the other hand, we did not find any significantly enriched upregulated GO terms. In cluster 3, we found that DEGs were enriched in upregulated GO terms associated with neuron function, olfaction, cell motility, and immune system. DEGs in cluster 3 were found to be enriched in downregulated GO terms associated with DNA damage response, ageing, and retinoid metabolism and response.

The difference in expression profiles illustrate highly heterogeneous transcriptomics in AD and PD and that there are notable commonalities and differences between the subclasses of AD or PD samples. Interestingly, we found retinoid metabolism or function to be a common altered GO term in all subclasses. This was upregulated in cluster 1 but downregulated in clusters 2 and 3. We therefore observed that retinoid dysregulation appears to be a common ageing-related hallmark of NDD.

### Metabolic analysis reveals retinoids and sex hormones as significantly dysregulated in AD and PD

Based on clustering and GSE analysis, we identified distinct expression profiles but these alone could not offer insights into metabolic activities of brain in AD and PD. To determine metabolic changes in the clusters compared to controls, we performed constraint-based genome-scale metabolic modelling. We reconstructed a brain-specific genome-scale metabolic model (GEM) based on the well-studied HMR2.0 (Mardinoglu et al., 2013) reference GEM by overlaying transcriptomic data from each cluster and applying brain-specific constraints as described previously (Baloni et al., 2020) using the tINIT algorithm (Agren et al., 2012, 2014) within the RAVEN Toolbox 2.0 (Wang et al., 2018). We generated a brain-specific GEM (*iBrain2845*) (**Supplementary file 1**) and used it as the reference GEM for reconstruction of cluster-specific GEMs in turn. We constructed the resulting context-specific *iADPD* series GEMs *iADPD1*, *iADPD2*, *iADPD3*, and *iADPDControl*, corresponding to cluster 1, cluster 2, cluster 3, and the control cluster, respectively (**Supplementary file 2**).

We conducted flux balance analysis (FBA) by defining maximisation of ATP synthesis as the objective function. *iADPD1* and *iADPD2* both showed upregulation of fluxes in reactions involved in cholesterol biosynthesis and downregulation in O-glycan metabolism, with reaction flux changes being more pronounced in *iADPD2* than in *iADPD1* (**Table 2**, **Supplementary data 2**). We found that the fluxes in *iADPD1* were uniquely upregulated in oestrogen metabolism and the Kandustch-Russell pathway. *iADPD2* was uniquely upregulated in cholesterol metabolism, whereas *iADPD3* uniquely displayed roughly equal parts upregulation and downregulation in several pathways, including aminoacyl-tRNA biosynthesis, androgen metabolism, arginine and proline metabolism, cholesterol biosynthesis, galactose metabolism, glycine, serine, and threonine metabolism, and N-glycan metabolism.

**Table 2.** Flux balance analysis of iADPD1, iADPD2, and iADPD3 versus iADPDControl. Flux balance analysis was performed for each *iADPD*-series GEM and the predicted fluxes for the three disease cluster GEMs were compared against the predicted fluxes for the control cluster GEM. Reactions are grouped by subsystem and flux difference values are expressed as mean flux difference between disease clusters and the control cluster across all changed reactions within a subsystem. For full results, refer to Supplementary data 2.

| **Subsystem** | ***iADPD1*** | ***iADPD2*** | ***iADPD3*** |
| --- | --- | --- | --- |
| Acyl-CoA hydrolysis | -0.001 | 0.001 | 0.000 |
| Alanine, aspartate and glutamate metabolism | -0.148 | 0.014 | 0.000 |
| Aminoacyl-tRNA biosynthesis | 4.698 | 4.698 | 0.000 |
| Androgen metabolism | -1.426 | -0.399 | -0.001 |
| Arachidonic acid metabolism | -0.098 | 0.010 | 0.000 |
| Arginine and proline metabolism | -0.182 | -0.327 | 0.000 |
| Beta oxidation of branched-chain fatty acids (mitochondrial) | -0.049 | -0.049 | -0.049 |
| Beta oxidation of di-unsaturated fatty acids (n-6) (mitochondrial) | -0.636 | 0.002 | -0.001 |
| Beta oxidation of odd-chain fatty acids (mitochondrial) | 0.001 | -0.002 | -0.002 |
| Beta oxidation of poly-unsaturated fatty acids (mitochondrial) | 0.709 | 0.024 | 0.000 |
| Beta oxidation of unsaturated fatty acids (n-7) (mitochondrial) | -0.016 | 0.001 | -0.003 |
| Beta oxidation of unsaturated fatty acids (n-9) (mitochondrial) | 0.011 | 0.000 | 0.007 |
| Carnitine shuttle (cytosolic) | 0.012 | 0.000 | -0.001 |
| Carnitine shuttle (mitochondrial) | 0.003 | 0.000 | 0.002 |
| Cholesterol biosynthesis 1 (Bloch pathway) | 0.076 | -0.983 | 0.001 |
| Cholesterol biosynthesis 2 | 2.501 | 4.472 | 0.000 |
| Cholesterol biosynthesis 3 (Kandustch-Russell pathway) | 1.699 | 0.000 | 0.000 |
| Cholesterol metabolism | 0.067 | 4.482 | 0.000 |
| Estrogen metabolism | 2.085 | 0.000 | 0.000 |
| Fatty acid activation (endoplasmic reticular) | 0.000 | 0.000 | 0.000 |
| Fatty acid biosynthesis (even-chain) | 0.000 | 0.000 | 0.000 |
| Fatty acid desaturation (even-chain) | 0.785 | 0.000 | 0.000 |
| Fatty acid elongation (odd-chain) | -0.042 | -0.024 | 0.000 |
| Formation and hydrolysis of cholesterol esters | -0.382 | 0.004 | 0.000 |
| Fructose and mannose metabolism | -0.211 | -0.007 | 0.000 |
| Galactose metabolism | -0.008 | 0.035 | 0.000 |
| Glycine, serine and threonine metabolism | 0.276 | 0.557 | 0.000 |
| Glycolysis / Gluconeogenesis | -0.213 | 0.022 | 0.033 |
| Histidine metabolism | 0.000 | 0.000 | 0.000 |
| Leukotriene metabolism | -0.032 | 0.000 | 0.000 |
| Lysine metabolism | 0.000 | 0.000 | 0.000 |
| N-glycan metabolism | -0.784 | 0.016 | 0.000 |
| Nitrogen metabolism | 0.000 | 0.000 | 0.000 |
| Nucleotide metabolism | 0.027 | -0.028 | 0.000 |
| O-glycan metabolism | -2.346 | -4.738 | 0.000 |
| Pentose phosphate pathway | 0.127 | 0.000 | 0.000 |
| Propanoate metabolism | -0.116 | 0.020 | 0.091 |
| Protein degradation | 0.000 | 0.000 | 0.000 |
| Purine metabolism | 0.112 | -0.013 | 0.000 |
| Pyrimidine metabolism | -0.071 | -0.010 | -0.001 |
| Pyruvate metabolism | -0.183 | -0.004 | -0.077 |
| Starch and sucrose metabolism | 0.000 | 0.000 | 0.000 |
| Steroid metabolism | -0.097 | -0.295 | 0.003 |
| Terpenoid backbone biosynthesis | 0.398 | 0.187 | 0.020 |
| Valine, leucine and isoleucine degradation | 0.127 | 0.000 | 0.000 |

In particular, we observed increased positive fluxes through reactions HMR_2055 and HMR_2059 in *iADPD1*, which convert oestrone to 2-hydroxyoestrone and then to 2-methoxyoestrone (**Figure 3**). In *iADPDControl*, these reactions carried zero flux. In *iADPD2*, we observed increased positive fluxes through HMR_1457 and HMR_1533, which produce geranyl pyrophosphate and lathosterol, respectively. Both of these molecules are precursors to cholesterol, and while we did not see a proportionate increase in the production of other molecules along the pathway (namely, farnesyl pyrophosphate and squalene), we did observe a general increase in fluxes through the androgen biosynthesis and metabolism pathway. Finally, we observed that *iADPD3* displayed a decreased production of testosterone from 4-androstene-3,17-dione via HMR_1974 despite an increase in production of 4-androstene-3,17-dione via HMR_1971.

**Figure 3.**
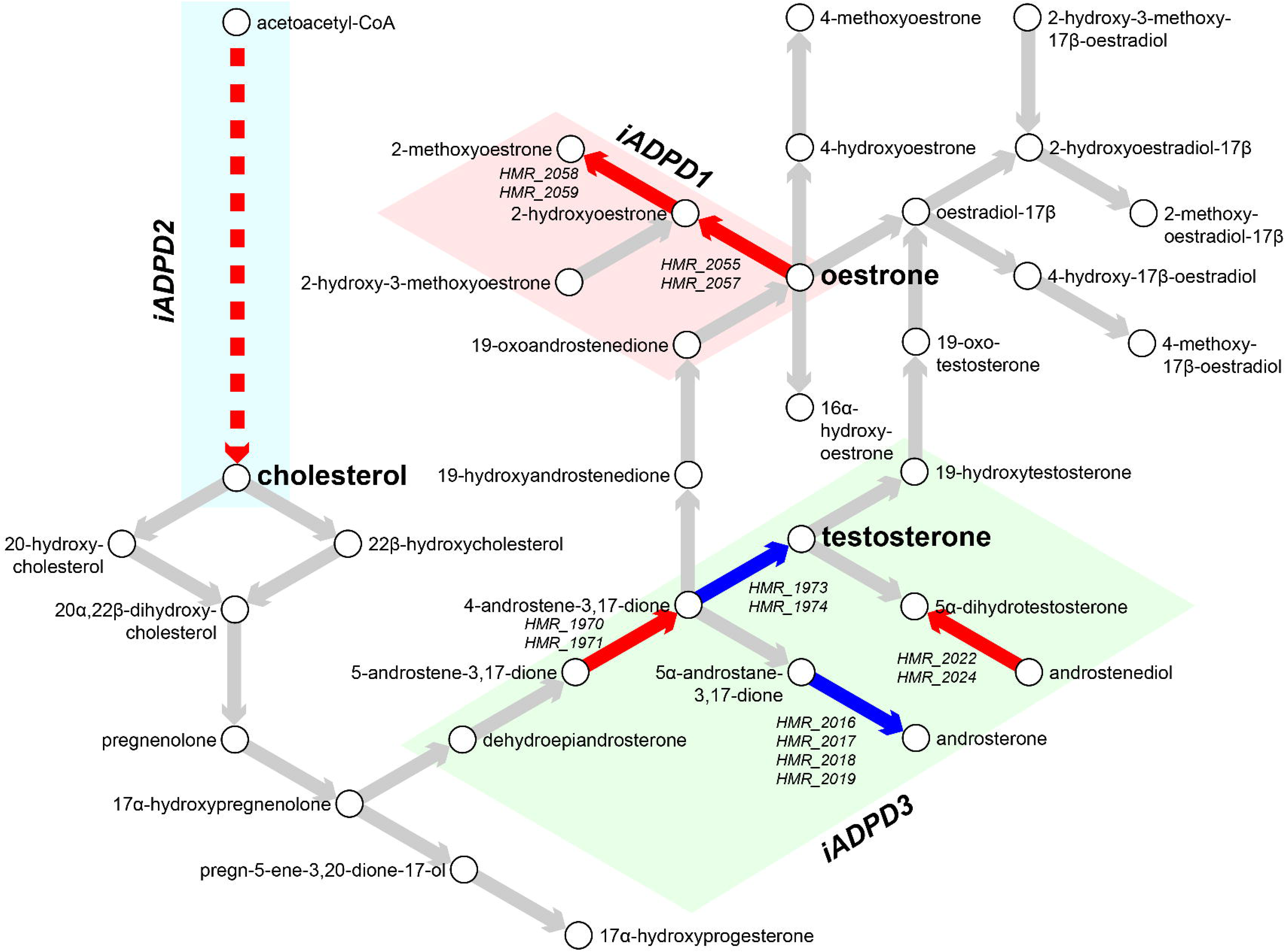
Metabolic characterisation of AD and PD subclasses. Flux balance analysis (FBA) was performed on *iADPD1-3* genome-scale metabolic models (GEMs) and flux values were compared with those of *iADPDControl*. Key metabolites and reactions within the androgen metabolism pathway are shown and key dysregulations are displayed as coloured arrows: red indicates increased flux compared to *iADPDControl*; blue indicates decreased flux compared to *iADPDControl*. Dysregulations associated to each GEM are shown in coloured boxes. The dashed line indicates multiple reactions are involved. Human Metabolic Reactions (HMR) identifiers are shown for androgen metabolism reactions with dysregulated fluxes. For full data, refer to Supplementary data 2.

Taken together, the obtained results indicate the existence of three distinct metabolic dysregulation profiles in AD and PD, with dysregulation being most pronounced in cluster 2 patients and least pronounced in cluster 3 patients. Furthermore, we found that all three feature dysregulations in or associated with sex hormone biosynthesis and metabolism, which might explain the heterogeneity in responses to sex hormone replacement therapy in AD and PD patients as extensively reported previously (Baum, 2005; Wahjoepramono et al., 2016; Resnick et al., 2017; Rajsombath et al., 2019). We also confirmed that dysregulations through sex hormone pathways in the *iADPD* series GEMs were not due to differences in relative frequencies between sexes in the main clusters 1-3 (Fisher’s exact test, *p* = 0.4700).

In addition to metabolic inference and FBA, we performed reporter metabolite analysis (Patil and Nielsen, 2005) by overlaying DEG analysis results onto the reference GEM to identify hotspots of metabolism (**Table 3**, **Supplementary data 3**). In short, we uniquely identified oestrone as a reporter metabolite in cluster 1, and lipids such as acylglycerol and dolichol in cluster 2. No notable reporter metabolites were identified as significantly changed in cluster 3 only. In common to all clusters 1-3, retinoids and sex hormones such as androsterone and pregnanediol were identified as significantly changed reporter metabolites, which are generally in line with GSE and FBA results.

**Table 3.** Reporter metabolite analysis of AD and PD subclasses. Reporter metabolite analysis was performed for each AD/PD subclass by overlaying differential expression results onto *iBrain2845*. Top 10 unique reporter metabolites by *p*-value for each cluster compared to the control cluster are shown. For full results, refer to Supplementary data 3.

| Reporter metabolite | Z-score | p-value |
| --- | --- | --- |
| **Cluster 1** |  |  |
| O2 | 6.111 | 4.95E-10 |
| estrone | 5.4557 | 2.44E-08 |
| retinoate | 5.3943 | 3.44E-08 |
| NADP+ | 5.3667 | 4.01E-08 |
| arachidonate | 5.2822 | 6.38E-08 |
| 2-hydroxyestradiol-17beta | 5.0999 | 1.70E-07 |
| linoleate | 5.0622 | 2.07E-07 |
| 10-HETE | 5.0454 | 2.26E-07 |
| 11,12,15-THETA | 5.0454 | 2.26E-07 |
| 11,14,15-theta | 5.0454 | 2.26E-07 |
| **Cluster 2** |  |  |
| 1-acylglycerol-3P-LD-PC pool | 4.3322 | 7.38E-06 |
| acyl-CoA-LD-PI pool | 4.143 | 1.71E-05 |
| phosphatidate-CL pool | 4.0973 | 2.09E-05 |
| thymidine | 3.5852 | 0.00016843 |
| uridine | 3.5852 | 0.00016843 |
| prostaglandin D2 | 3.2144 | 0.00065348 |
| G10596 | 3.1354 | 0.0008581 |
| G10597 | 3.1354 | 0.0008581 |
| D-myo-inositol-1,4,5-trisphosphate | 2.9988 | 0.0013552 |
| dolichyl-phosphate | 2.9655 | 0.001511 |
| **Cluster 3** |  |  |
| D-myo-inositol-1,4,5-trisphosphate | 2.6543 | 0.0039734 |
| 13-cis-retinal | 2.6537 | 0.0039806 |
| heparan sulfate, precursor 9 | 2.5915 | 0.0047772 |
| sn-glycerol-3-phosphate | 2.578 | 0.0049682 |
| DHAP | 2.5353 | 0.0056173 |
| porphobilinogen | 2.4987 | 0.0062333 |
| ATP | 2.4838 | 0.0064998 |
| L-glutamate 5-semialdehyde | 2.4576 | 0.006994 |
| prostaglandin D2 | 2.451 | 0.0071221 |
| ribose | 2.4133 | 0.0079045 |

### Network analysis supports retinoid and androgen dysregulation and suggests transcriptomic similarity between AD and PD

To further explore the gene expression patterns shown across AD and PD patients, we took expression data and constructed a weighted gene co-expression network for both groups (Spearman ρ > 0.9, FDR < 10^-9^, Materials and Methods). Each network was compared against equivalent randomly-generated networks as null models. After quality control, the AD network contained 4861 nodes (genes) and ∼397,000 edges (significant correlations), and the PD network contained 5857 nodes and ∼394,000 edges (**Figure 4A, Figure 4B**, **Table 4**). A community analysis to identify modules of highly co-expressed genes (Traag et al., 2019) highlighted nine and fifteen communities with significant functional enrichment in AD and PD respectively.

**Figure 4.**
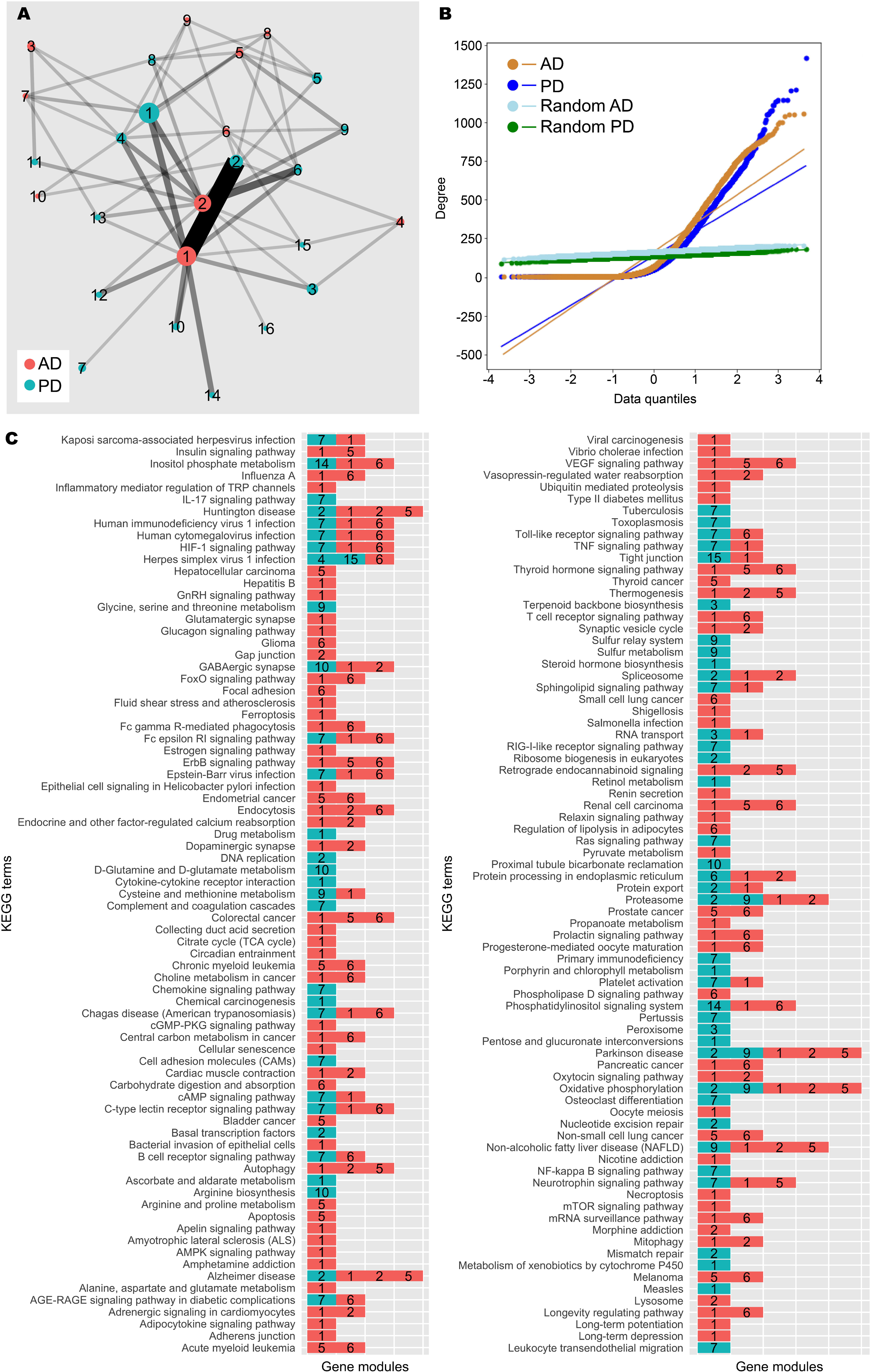
Network analysis of AD and PD gene co-expression modules. **A)** Gene co-expression networks were constructed from transcriptomic data from AD and PD samples. Community analysis was used to identify gene modules (Methods). Modules with at least 30 genes are shown as nodes. Node size indicates number of genes. Nodes are coloured by network of origin and numbered in descending order of module size. Shared genes between modules are shown as edges. Edge weight indicates number of shared genes. **B**) Degree distribution of AD, PD, and random networks. **C)** Enrichment analysis was performed on gene modules containing at least 30 genes using the KEGG database (Methods). Significantly enriched gene modules are shown as coloured, numbered blocks. Colour and number keys are as in (A).

**Table 4.** AD and PD network properties. Gene co-expression networks were generated for AD and PD samples. AD, PD, and random networks are shown.

|  | **Nodes** | **Edges** | **Diameter** | **Average path length** | **Density** | **Clustering coefficient** | **Connected network?** | **Minimum cut** |
| --- | --- | --- | --- | --- | --- | --- | --- | --- |
| **AD** | 4861 | 396985 | 11 | 3.004 | 0.034 | 0.443 | No | - |
| **PD** | 5857 | 394405 | 18 | 3.598 | 0.023 | 0.397 | No | - |
| **Random AD** | 4861 | 396985 | 3 | 1.970 | 0.034 | 0.034 | Yes | 114 |
| **Random PD** | 5857 | 394405 | 3 | 2.021 | 0.023 | 0.023 | Yes | 89 |

In the AD network, gene module C3 was enriched for genes involved with neuron and synapse development, similar to patient cluster 3, C4 for genes involved with mRNA splicing, similar to patient cluster 2, and C5 for genes involved with the mitochondrial electron transport chain (**Figure 4C**, **Supplementary data 4**). C1 and C2 were the gene modules with the largest number of genes. C1 was enriched for gene expression quality control genes and development and morphogenesis genes, mirroring patient cluster 2, whereas C2 contained cytoskeleton-related genes, similar to patient cluster 1.

In the PD network, C1 was enriched for genes involved with retinoid metabolism, glucuronidation, and cytokine signalling. Since androgens are major targets of glucuronidation (Grosse et al., 2013), these results are in line with our main findings. Further, C2 contained DNA damage response and gene regulation genes, similar to patient cluster 2, C3 contained nuclear protein regulation genes, and C4 contained mRNA splicing genes, again similar to patient cluster 2.

Further, the two networks share a large number of enriched terms in common, and there is high similarity between the major gene modules, highlighting the similarity between AD and PD. In addition to this, enrichment analysis for KEGG terms was unable to assign “Alzheimer disease” and “Parkinson disease” to the correct gene modules from the respective networks, and additional neurological disease terms such as “Huntington disease” and “Amyotrophic lateral sclerosis” were also identified by the analysis, further suggesting the transcriptomic similarity between neurological diseases. We found that AD C1 and PD C2 were frequently annotated with these disease terms, and these gene modules are also highly similar. Therefore, this gene module could constitute a core set of dysregulated genes in neurodegeneration.

Taken together, the network analysis supports our GSE findings. The functional consequences of differential expression in the patient clusters could be explained by differential modulation of gene modules identified in our network analysis together with dysregulation of a core set of genes implicated in both AD and PD.

### Zebrafish transcriptomic and metabolic investigations suggest an association between brain ageing and retinoid dysregulation

To further validate our findings regarding the differences between clusters of human AD and PD samples, we analysed transcriptomic data from *tert* mutant zebrafish and reconstructed tissue-specific GEMs (**Figure 5A**). To ascertain that these effects of ageing were limited to the brain, we analysed the brain, liver, muscle, and skin of zebrafish as well as the whole animal.

**Figure 5.**
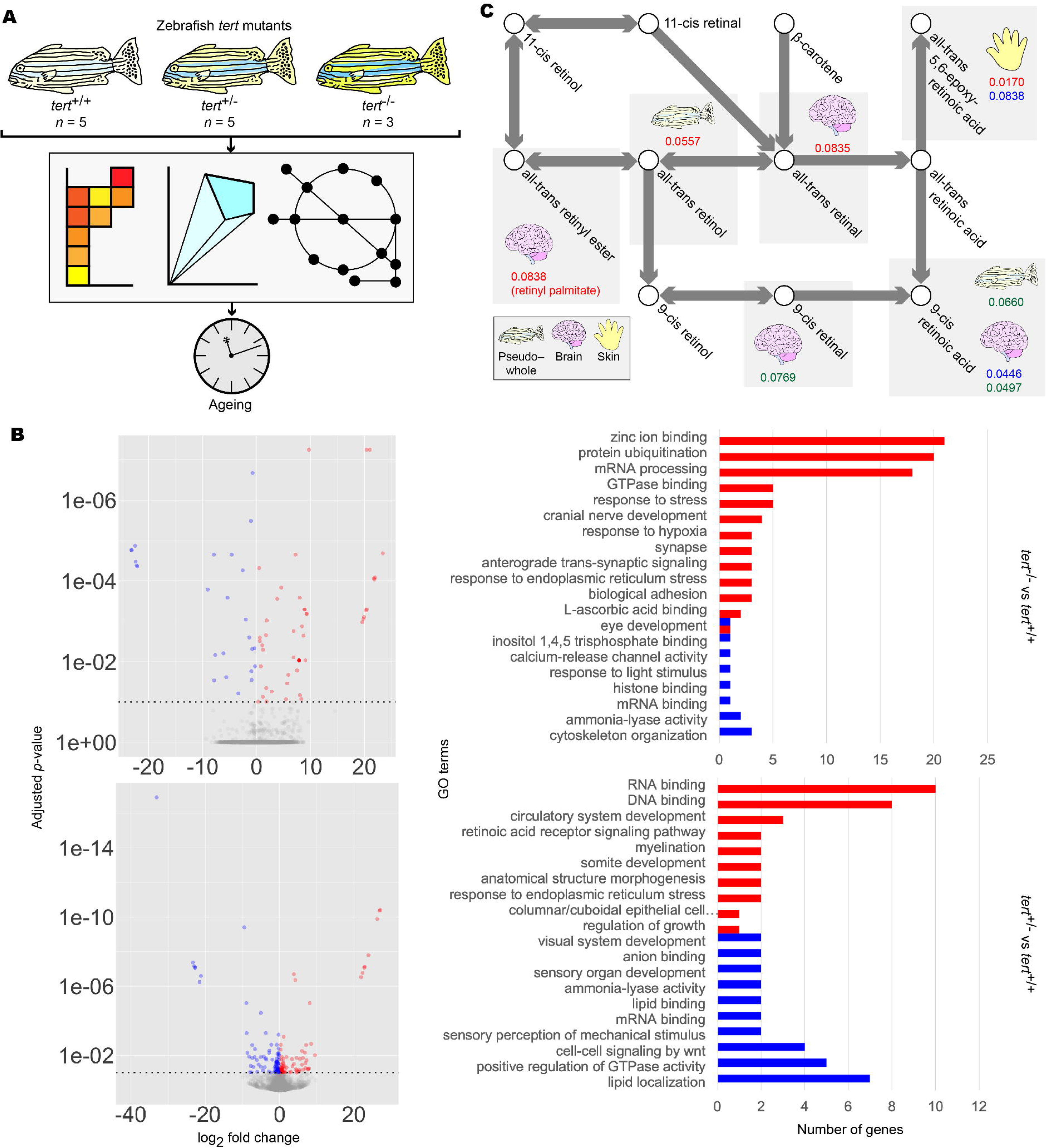
Summary of zebrafish *tert* mutant analysis. **A)** Workflow for the analysis of zebrafish *tert* mutants. **B)** Differentially expressed gene (DEG) (left panels) and gene set enrichment (GSE) analysis (right panels) of zebrafish brain samples. DEG and GSE analyses were performed on zebrafish *tert* mutant brain expression data for *tert*^-/-^ (upper panels) and *tert*^+/-^ (lower panels), using *tert*^+/+^ as a reference. Methods and colour keys are as in Figure 2. For muscle, liver, skin, and pseudo–whole animal analyses, refer to Supplementary figure 3. For full data, refer to Supplementary data 5. **C)** Reporter metabolite analysis of zebrafish samples. DEG data were overlaid on *ZebraGEM2.1* to determine reporter metabolites. Shown are reporter metabolites with *p* < 0.1 within the retinoic acid metabolic pathway. Red numbers indicate *p*-values in *tert*^-/-^ compared to *tert*^+/+^. Blue numbers indicate *p*-values in *tert*^+/-^ compared to *tert*^+/+^. Green numbers indicate *p*-values in *tert*^-/-^ compared to *tert*^+/-^. Tissues are indicated with icons. For full data, refer to Supplementary data 6.

We first repeated DEG and GSE analyses in the *tert* mutants using brain transcriptomic data. We found significant enrichment of GO terms associated with retinoid metabolism as well as eye development and light sensing, in which retinoids act as signalling molecules (Blomhoff and Blomhoff, 2006) (**Figure 5B**, **Supplementary figure 3, Supplementary data 5**). To further support our findings, we then reconstructed mutant- and genotype-specific GEMs by overlaying zebrafish *tert* mutant transcriptomic data onto a modified generic *ZebraGEM2* GEM (Van Steijn et al., 2019). We designated the modified GEM *ZebraGEM2.1* (**Supplementary file 3**) and used it as the reference GEM. We also generated zebrafish organ-specific GEMs and provide them to the interested reader (**Supplementary file 4**).

We then repeated reporter metabolite analysis using the transcriptomic data from zebrafish tissue-specific GEMs and found that retinoids were identified as significant reporter metabolites in *tert*^+/-^ zebrafish (*p* = 0.045) but not in *tert*^-/-^, where evidence was marginal (*p* = 0.084) (**Figure 5C**, **Table 5**, **Supplementary data 6**). We also observed this result in the skin of *tert*^-/-^ mutants, where evidence was significant (*p* = 0.017). This result can be explained due to the susceptibility of skin as an organ to photoageing, for which topical application of retinol is a widely-used treatment (Riahi et al., 2016). However, we did not find evidence for significant changes in pregnanediol, and androsterone was significant only in the skin of *tert*^-/-^ zebrafish (*p* = 0.017). This would suggest that either change in sex hormones are not ageing-related with regards AD and PD, or the changes were outside the scope of the zebrafish model that we used.

**Table 5.**
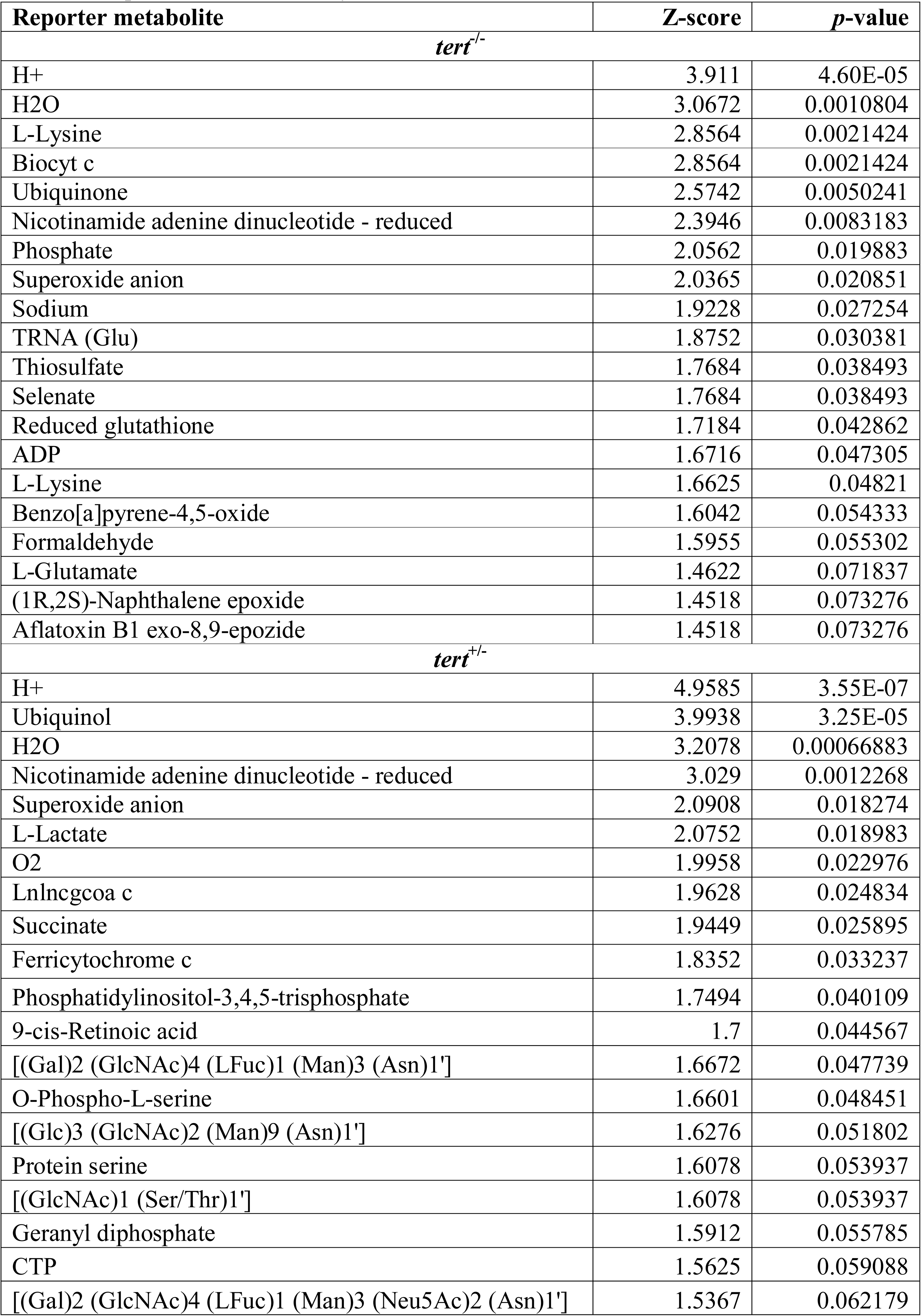
Reporter metabolite analysis of zebrafish *tert* mutants. Reporter metabolite analysis was performed for the brains of zebrafish *tert* mutant by overlaying differential expression results onto *ZebraGEM2.1*. Top 20 unique reporter metabolites by *p*-value for each cluster compared to wildtype *tert*^+/+^ zebrafish are shown. For full results, refer to Supplementary data 6.

Taken together, these results indicated that ageing can largely explain alterations in retinoid metabolism in the brain but not alterations in sex hormone metabolism. These results also suggested that ageing has a differential effect on different organs, implying that metabolic changes due to ageing in the brain are associated with neurological disorders.

## Discussion

In this work, we integrated gene expression data across diverse sources into context-specific GEMs and sought to identify and characterise disease subclasses of AD and PD. We used unsupervised clustering to identify AD/PD subclasses and employed DEG and GSE analysis to functionally characterise them. We used network exploration, constraint-based metabolic modelling, and reporter metabolite analysis to characterise flux and metabolic perturbations within basal metabolic functions and pathways. We then leveraged expression data from zebrafish ageing mutants to validate our findings that these perturbations might be explained by ageing. Our analysis concluded with the identification and characterisation of three AD/PD subclasses, each with distinct functional characteristics and metabolic profiles. All three subclasses showed depletion of retinoids by an ageing-related mechanism as a common characteristic.

We believe that a combined analysis that integrates AD and PD data is necessary to elucidate common attributes between the two diseases. However, we realised that such an analysis will likely obscure AD- and PD-specific factors, such as amyloid β and α-synuclein, but should aid the discovery of any factors in common. Since AD and PD share numerous risk factors and comorbidities such as old age, diabetes, and cancer risk, we believe that an AD/PD combined analysis can identify factors in common to both diseases and prove valuable for the identification of treatment strategies which might be effective in the treatment of both diseases.

GSE analysis highlighted significant changes related to retinoid function or visual system function, in which retinol and retinal act as signalling molecules (Blomhoff and Blomhoff, 2006), in all clusters (**Figure 2**, **Supplementary data 1**). Together with the identification of multiple retinol derivatives as significant reporter metabolites in *iBrain2845* (**Table 3**, **Supplementary data 3**), we hypothesised that retinoids are a commonly dysregulated class of molecules in both AD and PD, and that this may be due to an ageing mechanism. Indeed, in our investigation with zebrafish telomerase mutants, we again found alterations in retinoid and visual system function in GSE analysis (**Figure 5B**, **Supplementary figure 3**, **Supplementary data 5**) and reporter metabolite analysis (**Figure 5C**, **Table 5**, **Supplementary data 6**).

Retinoids were identified as a reporter metabolite in all three clusters of patients in this study, and we believe that retinoid therapy is a potentially viable treatment for both AD and PD patients. Further, our zebrafish analysis highlighted the importance of retinoids in ageing of the brain and the skin (**Figure 5C**, **Table 5**, **Supplementary data 6**). Retinol, its derivatives, and its analogues are already used as topical anti-ageing therapies for aged skin (Riahi et al., 2016), and there is a growing body of evidence suggesting its efficacy for the treatment of AD (Shudo et al., 2009; Fukasawa et al., 2012; Das et al., 2019; Fitz et al., 2019). We add to the body of evidence with this *in silico* investigation involving zebrafish telomerase mutants, suggesting that the source of retinoid depletion in AD and PD is ageing-related. Interestingly, regarding our finding for skin ageing in zebrafish, lipid biomarkers have been proposed in a recent skin sebum metabolomics study in PD patients (Sinclair et al., 2021). This could be interpreted as co-ageing in brain and skin tissues, possibly allowing for cheap, non-invasive prognostic testing for PD.

In addition to retinoids, we found evidence for subclass-specific dysregulation within the androgen metabolism pathway in each of the three clusters in FBA (**Table 2**, **Supplementary data 2**) and reporter metabolite analysis (**Table 3**, **Supplementary data 3**). We found that *iADPD1* displayed increased oestrone conversion to the less potent (Martucci, 1983) 2-methoxyoestrone, *iADPD2* displayed increased production of the cholesterol precursor molecules geranyl pyrophosphate and lathosterol and increased androgen biosynthesis, and *iADPD3* displayed decreased conversion of 4-androstene-3,17-dione to testosterone. However, there was no definitive evidence to suggest an ageing-related basis for these observations based on our zebrafish study, but this may be due to the diverse functional roles that sex hormones have, limitations within the *ZebraGEM2.1* model, or absence of an actual biological link between sex hormones and ageing of the brain. Despite this, given the widely reported variability in responses to sex hormone replacement therapy in AD and PD (Baum, 2005; Shepardson et al., 2011; Wahjoepramono et al., 2016; Resnick et al., 2017), we believe that this observation represents a possible explanation for the heterogeneity. Our observation regarding the dysregulation of the androgen pathway at three separate points suggests that dysregulation at other points might also be linked to AD and PD, thus implying that androgen metabolism dysregulation in general might be important for the development of AD and PD. Our finding via network community analysis of a gene module associated with glucuronidation activity points to a possible therapeutic strategy to combat androgen dysregulation. More work is needed to elucidate the importance of sex hormones and glucuronidation regarding AD and PD.

Identification of subclasses is desirable to address the heterogeneity in disease with regards transcriptomic profile and treatment response, but patients must be stratified in order to be diagnosed with the correct disease subclass and therefore administer the appropriate treatment. To this end, we used GSE analysis to functionally characterise the AD/PD subclasses (**Figure 2**, **Supplementary data 1**). Cluster 2, which was associated with a decreased immune and stress response, appeared to be most severe disease subclass, whereas cluster 3, which was associated with an increased sensory perception of smell, reduced haemostasis, and reduced immune and DNA damage response, seemed to be the least severe. Meanwhile, cluster 1 was associated with an increased immune and inflammatory responses and reduced sensory perception of smell. The functional terms are supported by community analysis of our AD and PD gene co-expression networks, which identified gene modules that roughly align with the GSE results (**Figure 4**, **Supplementary data 4**). The proposed severity ratings are supported by FBA findings, which show *iADPD2* as having the highest total flux dysregulation compared to control, and *iADPD3* as having the least (**Table 2**, **Supplementary data 2**). Although we did not attempt to characterise for stratifying and diagnosing patients in our study, our findings clearly showed that such stratification is possible. Given the differing nature of the proposed therapeutic strategies that we outline above, stratification of patients into distinct disease subclasses is desirable.

In conclusion, we report three distinct subclasses of AD and PD. The first subclass was identified as being associated with increased immune response, inflammatory response, and reduced sensory perception of smell, according to GSE results. We observed that this subclass exhibited increased oestradiol turnover, according to FBA results. We therefore propose that subjects consistent with the first subclass may be treatable with combined retinoid and oestradiol therapy. The second subclass was linked with increased cholesterol biosynthesis and general increased flux through the androgen biosynthesis and metabolism pathway. This subclass was characterised by reduced immune response. We therefore suggest that subjects consistent with the second subclass be studied further with combined retinoid and statin therapy. The third subclass was characterised by enrichment of GO terms indicating increased sensory perception of smell, reduced haemostasis, and reduced immune and DNA damage response. This subclass also exhibited reduced testosterone biosynthesis from androstenedione, as determined by FBA. We therefore hypothesise that subjects consistent with the third subclass may benefit from combined retinoid and testosterone therapy. For all subclasses of AD and PD, more investigation is required to verify the effectiveness of these stratification methods and precision therapies. To our knowledge this is the first meta-analysis at this scale highlighting the potential significance of NDD therapy using retinoids, oestradiol, and testosterone by studying AD and PD in combination. We observed that the existence of disease subclasses demands precision or personalised medicine and explains the heterogeneity in NDD response to single-factor treatments.

## Materials and methods

### Data acquisition and processing

Gene expression values of protein-coding genes from the ROSMAP dataset were determined using kallisto (Bray et al., 2016) by aligning raw RNA sequencing reads to the *Homo sapiens* genome in Ensembl release 96 (Yates et al., 2020). Raw single-cell RNA sequencing reads from ROSMAP were converted to counts in Cell Ranger 4.0 (10X Genomics, https://support.10xgenomics.com/single-cell-gene-expression/software/pipelines/latest/installation) and aligned to the Cell Ranger *Homo sapiens* reference transcriptome version 2020-A. Single-cell expression values were compiled into pseudo-bulk expression profiles for each sample.

AD, PD, and control brain expression values of protein-coding genes from the ROSMAP dataset (Myers et al., 2007; Webster et al., 2009; Mostafavi et al., 2018), GTEx database version 8 (GTEx Consortium, 2013), FANTOM5 database (Forrest et al., 2014; Lizio et al., 2015, 2019) via Regulatory Circuits Network Compendium 1.0 (Marbach et al., 2016), HPA database (Uhlén et al., 2015), Rajkumar dataset (Rajkumar et al., 2020), and Zhang/Zheng dataset (Zhang et al., 2005; Zheng et al., 2010) were then combined. Genes from GTEx and FANTOM5 brain samples were filtered such that only genes whose products are known to participate in a protein-protein interaction described in the HuRI database (Luck et al., 2019) were included. Expression values were scaled and TMM normalised per sample, Pareto scaled per gene, and batch effects removed with the *removeBatchEffects* function from the limma (Ritchie et al., 2015) R package. After quality control and normalisation, a total of 64794 genes and 2055 samples resulted, of which 1572 samples corresponding to AD, PD, or control were accepted for analysis.

Projections onto 2-D space by PCA, t-SNE (Van Der Maaten and Hinton, 2008), and UMAP (McInnes et al., 2018) methods were generated on data after missing value imputation with data diffusion (van Dijk et al., 2018). t-SNE projections were generated with perplexity 20 and 1000 iterations. All other parameters were kept default. PCA and UMAP projections were generated using all default parameters.

### Transcriptome analysis

Using normalised, imputed expression values, AD and PD samples were then arranged into clusters without supervision using ConsensusClusterPlus (Wilkerson and Hayes, 2010) with maxK = 20 and rep = 1000. All other parameters were kept default. Clustering by *k* = 3 clusters was selected for downstream analysis. A fourth cluster containing only control samples was artificially added to the analysis.

For differential gene expression analysis, normalised, non-imputed counts were used. Genes were removed if expression values were missing in 40% or more of samples or were zero in all samples. Differential expression was then performed using DESeq2 (Love et al., 2014) with uniform size factors and all other parameters set to default. Genes with a Benjamini-Hochberg adjusted *p*-value at or below a cut-off of 1×10^-10^ were determined significantly differentially expressed genes.

Gene set enrichment analysis was performed using piano (Väremo et al., 2013) using all default parameters. GO term lists were obtained from Ensembl Biomart [https://www.ensembl.org/biomart/martview, accessed 2021-03-09] and were used as gene set collections. Enrichment of GO terms was determined by analysing GO terms of genes differentially expressed genes detected by DESeq2 as well as the parents of those GO terms. GO terms with an adjusted *p*-value at or below 0.05 for distinct-directional and/or mixed-directional methods were determined statistically significant.

### Metabolic analysis

For each cluster, consensus gene expression values were determined by taking the geometric mean of normalised expression counts across all samples within each cluster.

A reference GEM was created by modifying the gene associations of all reactions within the adipocyte-specific GEM *iAdipocytes1850* (Mardinoglu et al., 2013) to match those within the generic human GEM HMR3 (Mardinoglu et al., 2014). The resulting GEM was designated *iBrain2845*. Cluster-specific GEMs were reconstructed using the RAVEN Toolbox 2.0 (Wang et al., 2018) tINIT algorithm (Agren et al., 2012, 2014) with *iBrain2845* as the reference GEM.

FBA was conducted on each cluster-specific GEM using the *solveLP* function from the RAVEN Toolbox 2.0 with previously reported constraints (Baloni et al., 2020) and defining ATP synthesis (*iBrain2845:* HMR_6916) as the objective function. All constraints were applied with the exception of the following reaction IDs, which were excluded: EX_ac[e] (*iBrain2845*: HMR_9086) and EX_etoh[e] (*iBrain2845*: HMR_9099).

Reporter metabolite analysis was conducted using the *reporterMetabolites* function (Patil and Nielsen, 2005) from the RAVEN Toolbox 2.0, using *iBrain2845* as the reference model.

### Network analysis

To generate gene networks, normalised, non-imputed expression values from AD and PD samples were taken. Control samples and samples from blood were excluded. One network was generated each for AD and PD. For the AD model, all male samples were included and 171 female samples were chosen at random and included. For the PD model, all samples were included. Genes with any missing values were dropped. Genes with the 15% lowest expression or 15% lowest variance were disregarded from further analysis. Spearman correlations were calculated for each pair of genes and the top 1% of significant correlations were used to generate gene co-expression networks. Random Erdős-Rényi models were created for the AD and PD models with the same numbers of nodes and edges to act as null models, and compared against their respective networks in terms of centrality distributions. Community analyses were performed through the Leiden algorithm (Traag et al., 2019) by optimizing CPMVertexPartition, after a resolution scan of 10,000 points between 10^-3^ and 10. The scan showed global maxima at resolutions = 0.077526 and 0.089074 for AD and PD networks, which were used for optimization. Enrichment analysis was performed on modules with >30 nodes using enrichr (Chen et al., 2013; Kuleshov et al., 2016) using GO Biological Process, KEGG, and Online Mendelian Inheritance in Man libraries and was explored using Revigo (Supek et al., 2011).

### Zebrafish data acquisition and analysis

The *tert* mutant zebrafish line (*tert*^hu3430^) was obtained from Miguel Godhino Ferreira (Henriques et al., 2013). Fish maintenance, RNA isolation, processing, and sequencing were conducted as described previously (Aramillo Irizar et al., 2018).

From *n* = 5 wildtype (*tert*^+/+^), *n* = 5 heterozygous mutant (*tert*^+/-^), and *n* = 3 homozygous mutant (*tert*^-/-^), expression values were determined from RNA sequencing reads using kallisto by aligning to the *Danio rerio* genome in Ensembl release 96 (Yates et al., 2020). Expression values were generated for each extracted tissue as well as ‘psuedo–whole animal’, containing combined values across all tissues.

A reference zebrafish GEM was manually curated by modifying the existing *ZebraGEM2* model and was designated *ZebraGEM2.1*.

Differential expression analysis, gene set enrichment analysis, GEM reconstruction, FBA, and reporter metabolite analysis were conducted on *tert*^-/-^ and *tert*^+/-^ animals against a *tert*^+/+^ reference using DESeq2, piano, and RAVEN Toolbox 2.0 with default parameters. Reporter metabolite analysis was conducted with *ZebraGEM2.1* as the reference GEM.

FBA was attempted as described for the human GEMs with the exception that the following metabolic constraints were excluded: r1391, HMR_0482 (*ZebraGEM2.1*: G3PDm), EX_ile_L[e] (*ZebraGEM2.1*: EX_ile_e), EX_val_L[e] (*ZebraGEM2.1*: EX_val_e), EX_lys_L[e] (*ZebraGEM2.1*: EX_lys_e), EX_phe_L[e] (*ZebraGEM2.1*: EX_phe_e), GLCt1r, EX_thr_L[e] (*ZebraGEM2.1*: EX_thr_e), EX_met_L[e] (*ZebraGEM2.1*: EX_met L_e), EX_arg_L[e] (*ZebraGEM2.1*: EX_arg_e), EX_his_L[e] (*ZebraGEM2.1*: EX_his L_e), EX_leu_L[e] (*ZebraGEM2.1*: EX_leu_e), and EX_o2[e] (*ZebraGEM2.1*: EX_o2_e). The objective function was defined as ATP synthesis (*ZebraGEM2.*1: ATPS4m). FBA results for zebrafish are not presented.

### Data and code accessibility

All original computer code, models, and author-curated data files have been released under a Creative Commons Attribution ShareAlike 4.0 International Licence (https://creativecommons.org/licenses/by-sa/4.0/) and are freely available for download from <https://github.com/SimonLammmm/ad-pd-retinoid>.

Zebrafish *tert* mutant sequencing data have been deposited in the NCBI Gene Expression Omnibus (GEO) and are accessible through GEO Series accession numbers GSE102426, GSE102429, GSE102431, and GSE102434.

## Ethics statement

Zebrafish were housed in the fish facility of the Leibniz Institute on Aging – Fritz Lipmann Institute (FLI) under standard conditions and a 14-h–light and 10-h–dark cycle. All animal procedures were performed in accordance with the German animal welfare guidelines and approved by the Landesamt fu□r Verbraucherschutz Th□ringen (TLV), Germany.

## Author contributions

A.M. supervised the study and designed the study on human data. R.K. and C.E. designed the study on zebrafish. N.H. performed the zebrafish RNA sequencing and generated the raw counts. With the exception of the network analysis, S.L. performed all *in silico* analysis and M.A. and R.B. provided technical consultation. R.B. performed the network analysis. S.L. analysed all the results. S.L. wrote the manuscript with input from all the authors.

## Acknowledgements

We are grateful to Catarina Henriques and Miguel Godinho Ferreira for sharing the *tert* mutant zebrafish line. We also thank Ivonne Heinze, Ivonne Görlich, and Marco Groth from the FLI sequencing facility for sequencing the zebrafish samples.

The authors acknowledge use of the research computing facility at King’s College London, *Rosalind* (https://rosalind.kcl.ac.uk).

The Genotype-Tissue Expression (GTEx) Project was supported by the Common Fund of the Office of the Director of the National Institutes of Health, and by NCI, NHGRI, NHLBI, NIDA, NIMH, and NINDS. The data used for the analyses described in this manuscript were obtained from the GTEx Portal on 2019-12-06.

The results published here are in part based on data obtained from the AD Knowledge Portal (https://adknowledgeportal.synapse.org). Study data were provided by the Rush Alzheimer’s Disease Center, Rush University Medical Center, Chicago. Data collection was supported through funding by NIA grants P30AG10161 (ROS), R01AG15819 (ROSMAP; genomics and RNAseq), R01AG17917 (MAP), R01AG30146, R01AG36042 (5hC methylation, ATACseq), RC2AG036547 (H3K9Ac), R01AG36836 (RNAseq), R01AG48015 (monocyte RNAseq) RF1AG57473 (single nucleus RNAseq), U01AG32984 (genomic and whole exome sequencing), U01AG46152 (ROSMAP AMP-AD, targeted proteomics), U01AG46161(TMT proteomics), U01AG61356 (whole genome sequencing, targeted proteomics, ROSMAP AMP-AD), the Illinois Department of Public Health (ROSMAP), and the Translational Genomics Research Institute (genomic). Additional phenotypic data can be requested at www.radc.rush.edu. We thank the patients and their families for their selfless donation to further understanding Alzheimer’s disease. This project was supported by funding from the National Institute on Aging (AG034504 and AG041232). Many data and biomaterials were collected from several National Institute on Aging (NIA) and National Alzheimer’s Coordinating Center (NACC, grant #U01 AG016976) funded sites. Amanda J. Myers, PhD (University of Miami, Department of Psychiatry) prepared the series. The directors, pathologist and technicians involved include: Rush University Medical Center, Rush Alzheimer’s Disease Center (NIH #AG10161): David A. Bennett, M.D. Julie A. Schneider, MD, MS, Karen Skish, MS, PA (ASCP)MT, Wayne T Longman. The Rush portion of this study was supported by National Institutes of Health grants P30AG10161, R01AG15819, R01AG17917, R01AG36042, R01AG36836, U01AG46152, R01AG34374, R01NS78009, U18NS82140, R01AG42210, R01AG39478, and the Illinois Department of Public Health. - Quality control checks and preparation of the gene expression data was provided by the National Institute on Aging Alzheimer’s Disease Data Storage Site (NIAGADS, U24AG041689) at the University of Pennsylvania.

## Supplementary figure legends

**Supplementary figure 1.**
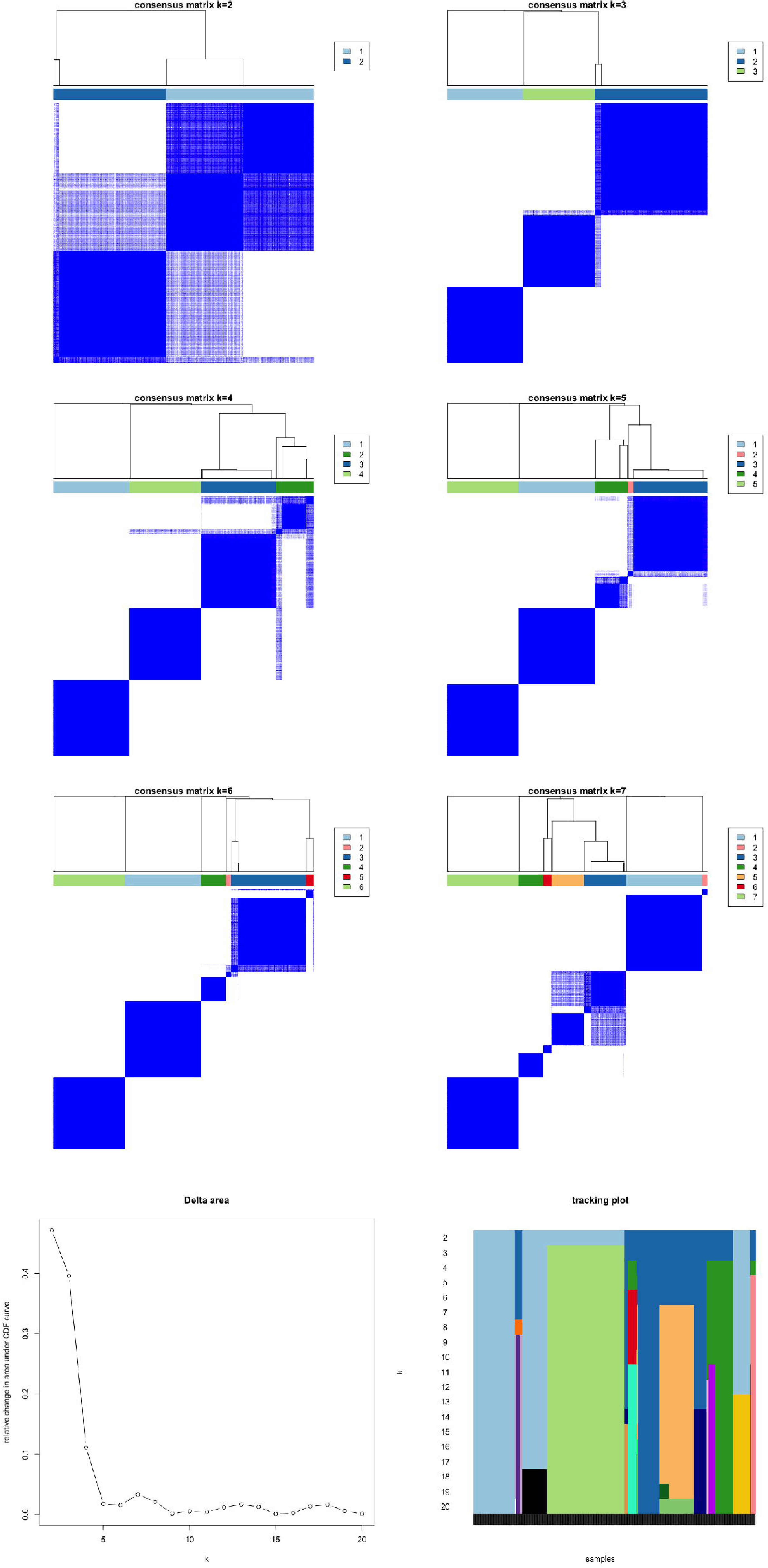
Unsupervised clustering of AD and PD samples. AD and PD samples were clustered into *k* clusters without supervision on the basis of normalised expression counts. Clustering was performed with 2 ≤ *k* ≤ 20. Consensus matrices for 2 ≤ *k* ≤ 7 are shown. Parameters and colour keys are as in Figure 1b.

**Supplementary figure 2.**
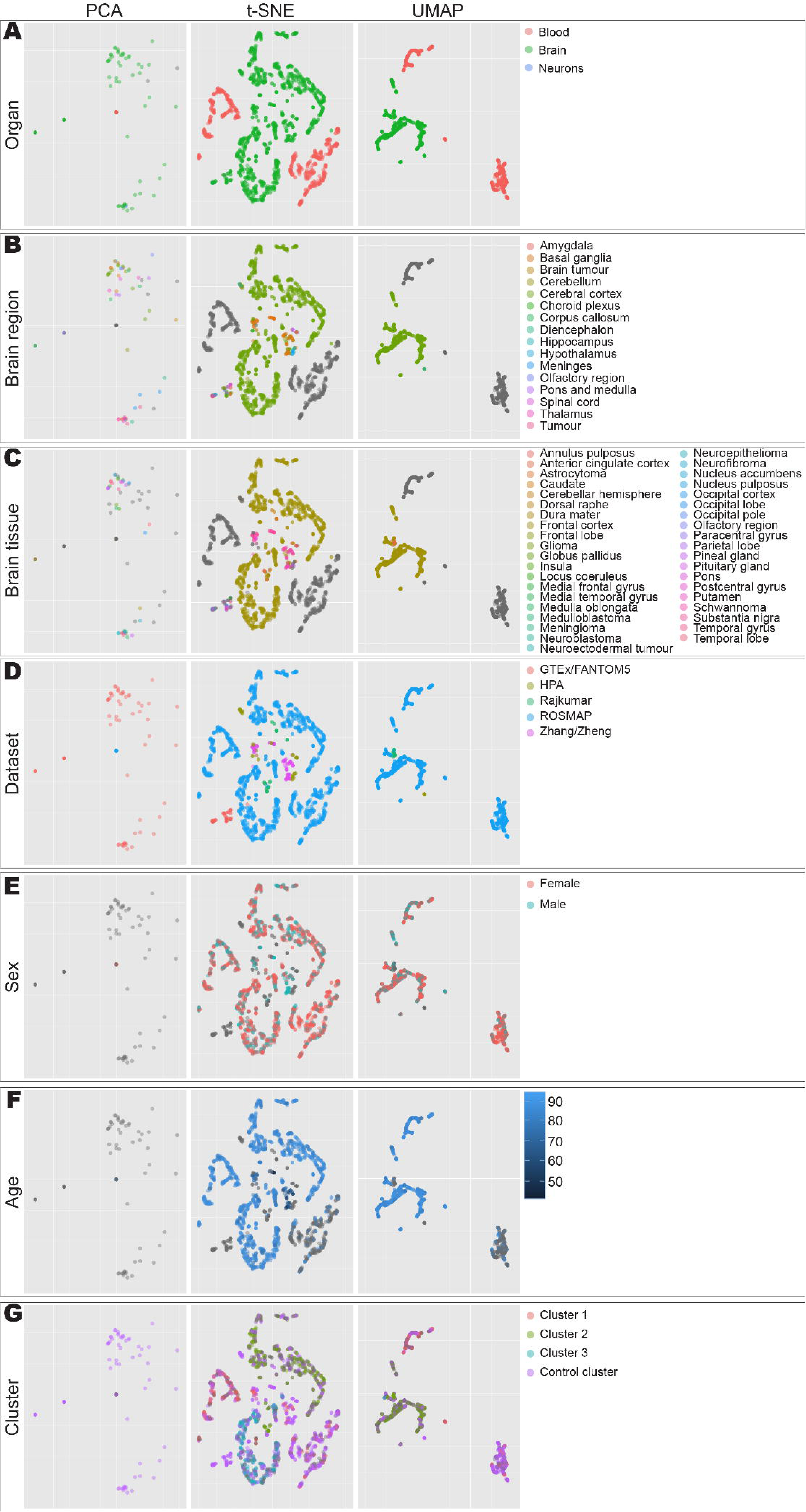
Visualisation of AD and PD samples. Expression data from AD and PD and control samples were integrated, normalised, and projected onto 2-D space using principal component analysis (PCA), t-distributed stochastic neighbour embedding (t-SNE), and uniform manifold approximation and projection (UMAP). Points are coloured according to **A)** organ of sample origin, **B)** brain subregion of sample origin, **C)** brain tissue of sample origin, **D)** dataset, **E)** sex, **F)** age, or **G)** cluster assignment by unsupervised clustering. Points with no data available are shown in grey.

**Supplementary figure 3.**
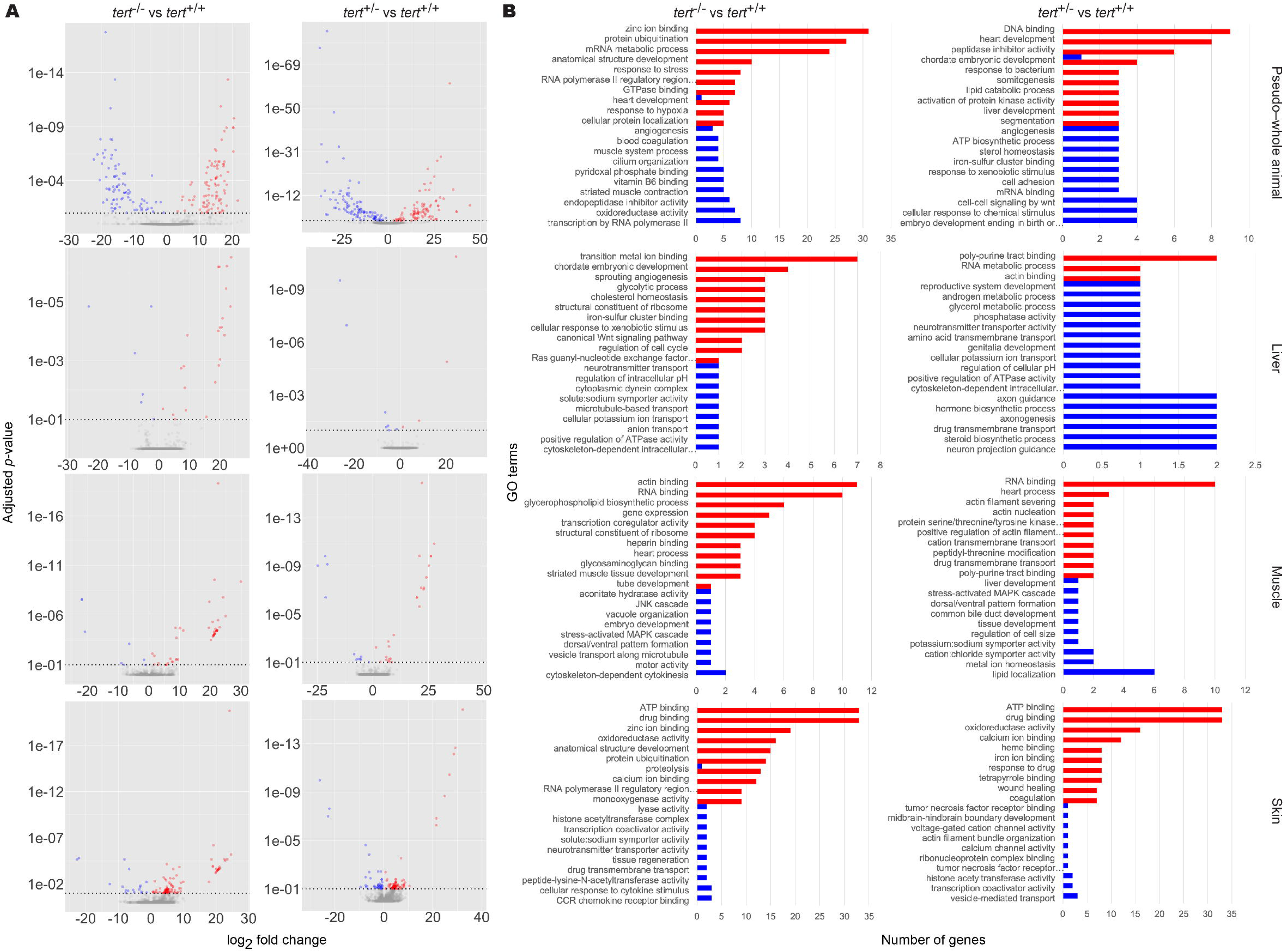
Transcriptomic and functional characterisation of zebrafish *tert* mutants. Differentially expressed gene (DEG) and gene set enrichment (GSE) analyses were performed on zebrafish *tert* mutant expression data for *tert^-/-^* and *tert^+/-^*, using *tert*^+/+^ as a reference. Methods and colour keys are as in Figure 2. **A)** DEG analyses. **B)** GSE analyses. Left panels, *tert*^-/-^ vs *tert*+/+. Right panels, *tert*^+/-^ vs *tert*^+/+^. Panels top to bottom: pseudo–whole animal, liver, muscle, skin. For the brain, refer to Figure 5. For full results, refer to Supplementary data 5.

## Supplementary files and data

Supplementary file 1. *iBrain2845* genome-scale metabolic model.

Supplementary file 2. *iADPD*-series context-specific genome-scale metabolic models.

Supplementary file 3. *ZebraGEM2.1* genome-scale metabolic model.

Supplementary file 4. Zebrafish context-specific genome-scale metabolic models.

Supplementary data 1. Gene set enrichment analysis results for AD and PD subclasses.

Supplementary data 2. Flux balance analysis results for *iADPD*-series genome-scale metabolic models.

Supplementary data 3. Reporter metabolite analysis results for AD and PD subclasses.

Supplementary data 4. Network analysis results for AD and PD samples.

Supplementary data 5. Gene set enrichment analysis results for zebrafish *tert* mutants.

Supplementary data 6. Reporter metabolite analysis results for zebrafish *tert* mutants.

## References

Agren R, Bordel S, Mardinoglu A, Pornputtapong N, Nookaew I, Nielsen J (2012) Reconstruction of genome-scale active metabolic networks for 69 human cell types and 16 cancer types using INIT Maranas CD, ed. PLoS Comput Biol 8:e1002518 Available at: http://dx.plos.org/10.1371/journal.pcbi.1002518 [Accessed January 15, 2020].

Agren R, Mardinoglu A, Asplund A, Kampf C, Uhlen M, Nielsen J (2014) Identification of anticancer drugs for hepatocellular carcinoma through personalized genome-scale metabolic modeling. Mol Syst Biol 10:721.

Altay O, Nielsen J, Uhlen M, Boren J, Mardinoglu A (2019) Systems biology perspective for studying the gut microbiota in human physiology and liver diseases. EBioMedicine 49:364–373 Available at: https://linkinghub.elsevier.com/retrieve/pii/S2352396419306486 [Accessed November 21, 2019].

Anchelin M, Alcaraz-Pérez F, Martínez CM, Bernabé-García M, Mulero V, Cayuela ML (2013) Premature aging in telomerase-deficient zebrafish. DMM Dis Model Mech 6:1101–1112 Available at: /pmc/articles/PMC3759330/?report=abstract [Accessed January 14, 2021].

Aramillo Irizar P et al. (2018) Transcriptomic alterations during ageing reflect the shift from cancer to degenerative diseases in the elderly. Nat Commun 9:327 Available at: https://pubmed.ncbi.nlm.nih.gov/29382830/ [Accessed February 15, 2021].

Bajaj A, Driver JA, Schernhammer ES (2010) Parkinson’s disease and cancer risk: A systematic review and meta-analysis. Cancer Causes Control 21:697–707.

Baloni P et al. (2020) Metabolic Network Analysis Reveals Altered Bile Acid Synthesis and Metabolism in Alzheimer’s Disease. Cell Reports Med 1:100138 Available at: https://doi.org/10.1016/j.xcrm.2020.100138 [Accessed January 19, 2021].

Baum LW (2005) Sex, hormones, and Alzheimer’s disease. Journals Gerontol - Ser A Biol Sci Med Sci 60:736–743 Available at: https://academic.oup.com/biomedgerontology/article-lookup/doi/10.1093/gerona/60.6.736 [Accessed January 21, 2021].

Blomhoff R, Blomhoff HK (2006) Overview of retinoid metabolism and function. J Neurobiol 66:606–630 Available at: https://pubmed.ncbi.nlm.nih.gov/16688755/ [Accessed February 17, 2021].

Bray NL, Pimentel H, Melsted P, Pachter L (2016) Near-optimal probabilistic RNA-seq quantification. Nat Biotechnol 34:525–527.

Carneiro MC, De Castro IP, Ferreira MG (2016) Telomeres in aging and disease: Lessons from zebrafish. DMM Dis Model Mech 9:737–748 Available at: https://dmm.biologists.org/content/9/7/737 [Accessed November 17, 2020].

Chakravarti D, LaBella KA, DePinho RA (2021) Telomeres: history, health, and hallmarks of aging. Cell 184:306–322 Available at: http://www.ncbi.nlm.nih.gov/pubmed/33450206 [Accessed January 25, 2021].

Chen EY, Tan CM, Kou Y, Duan Q, Wang Z, Meirelles G V., Clark NR, Ma’ayan A (2013) Enrichr: Interactive and collaborative HTML5 gene list enrichment analysis tool. BMC Bioinformatics 14:128 Available at: https://pubmed.ncbi.nlm.nih.gov/23586463/ [Accessed March 24, 2021].

Das B, Dasgupta S, Ray S (2019) Potential therapeutic roles of retinoids for prevention of neuroinflammation and neurodegeneration in Alzheimer’s disease. Neural Regen Res 14:1880–1892 Available at: https://pubmed.ncbi.nlm.nih.gov/31290437/ [Accessed January 21, 2021].

De La Monte SM, Wands JR (2008) Alzheimer’s disease is type 3 diabetes-evidence reviewed. J Diabetes Sci Technol 2:1101–1113.

Driver JA, Beiser A, Au R, Kreger BE, Splansky GL, Kurth T, Kiel DP, Lu KP, Seshadri S, Wolf PA (2012) Inverse association between cancer and Alzheimer’s disease: Results from the Framingham Heart Study. BMJ 344:e1442.

Fitz NF, Nam KN, Koldamova R, Lefterov I (2019) Therapeutic targeting of nuclear receptors, liver X and retinoid X receptors, for Alzheimer’s disease. Br J Pharmacol 176:3599–3610 Available at: https://pubmed.ncbi.nlm.nih.gov/30924124/ [Accessed January 21, 2021].

Forrest ARR et al. (2014) A promoter-level mammalian expression atlas. Nature 507:462–470.

Fukasawa H, Nakagomi M, Yamagata N, Katsuki H, Kawahara K, Kitaoka K, Miki T, Shudo K (2012) Tamibarotene: A candidate retinoid drug for Alzheimer’s disease. Biol Pharm Bull 35:1206–1212 Available at: https://pubmed.ncbi.nlm.nih.gov/22863914/ [Accessed January 21, 2021].

Greenland JC, Williams-Gray CH, Barker RA (2019) The clinical heterogeneity of Parkinson’s disease and its therapeutic implications. Eur J Neurosci 49:328–338 Available at: http://doi.wiley.com/10.1111/ejn.14094 [Accessed January 25, 2021].

Grosse L, Pâquet S, Caron P, Fazli L, Rennie PS, Bélanger A, Barbier O (2013) Androgen glucuronidation: An unexpected target for androgen deprivation therapy, with prognosis and diagnostic implications. Cancer Res 73:6963–6971 Available at: http://cancerres.aacrjournals.org/ [Accessed March 7, 2021].

GTEx Consortium (2013) The Genotype-Tissue Expression (GTEx) project. Nat Genet 45:580–585.

Henriques CM, Carneiro MC, Tenente IM, Jacinto A, Ferreira MG (2013) Telomerase Is Required for Zebrafish Lifespan. PLoS Genet 9:1003214 Available at: /pmc/articles/PMC3547866/ [Accessed February 15, 2021].

Hindle J V. (2010) Ageing, neurodegeneration and Parkinson’s disease. Age Ageing 39:156–161 Available at: https://academic.oup.com/ageing/article-lookup/doi/10.1093/ageing/afp223 [Accessed January 25, 2021].

Jeong SM, Jang W, Shin DW (2019) Association of statin use with Parkinson’s disease: Dose–response relationship. Mov Disord 34:1014–1021 Available at: https://onlinelibrary.wiley.com/doi/abs/10.1002/mds.27681 [Accessed January 25, 2021].

Joshi A, Rienks M, Theofilatos K, Mayr M (2020) Systems biology in cardiovascular disease: a multiomics approach. Nat Rev Cardiol 18:313–330 Available at: https://www.nature.com/articles/s41569-020-00477-1 [Accessed January 25, 2021].

Kuleshov M V., Jones MR, Rouillard AD, Fernandez NF, Duan Q, Wang Z, Koplev S, Jenkins SL, Jagodnik KM, Lachmann A, McDermott MG, Monteiro CD, Gundersen GW, Ma’ayan A (2016) Enrichr: a comprehensive gene set enrichment analysis web server 2016 update. Nucleic Acids Res 44:W90–W97 Available at: https://pubmed.ncbi.nlm.nih.gov/27141961/ [Accessed March 24, 2021].

Lam S, Bayraktar A, Zhang C, Turkez H, Nielsen J, Boren J, Shoaie S, Uhlen M, Mardinoglu A (2020) A systems biology approach for studying neurodegenerative diseases. Drug Discov Today 25:1146–1159 Available at: https://doi.org/10.1016/j.drudis.2020.05.010.

Lam S, Doran S, Yuksel HH, Altay O, Turkez H, Nielsen J, Boren J, Uhlen M, Mardinoglu A (2021) Addressing the heterogeneity in liver diseases using biological networks. Brief Bioinform 22:1751–1766 Available at: https://doi.org/10.1093/bib/bbaa002.

Liberini P, Valerio A, Memo M, Spano PF (1996) Lewy-body dementia and responsiveness to cholinesterase inhibitors: A paradigm for heterogeneity of Alzheimer’s disease? Trends Pharmacol Sci 17:155–160.

Lizio M et al. (2015) Gateways to the FANTOM5 promoter level mammalian expression atlas. Genome Biol 16:22 Available at: https://genomebiology.biomedcentral.com/articles/10.1186/s13059-014-0560-6 [Accessed January 14, 2021].

Lizio M, Abugessaisa I, Noguchi S, Kondo A, Hasegawa A, Hon CC, De Hoon M, Severin J, Oki S, Hayashizaki Y, Carninci P, Kasukawa T, Kawaji H (2019) Update of the FANTOM web resource: Expansion to provide additional transcriptome atlases. Nucleic Acids Res 47:D752–D758 Available at: http://fantom.gsc.riken. [Accessed January 14, 2021].

Long JM, Holtzman DM (2019) Alzheimer Disease: An Update on Pathobiology and Treatment Strategies. Cell 179:312–339 Available at: /pmc/articles/PMC6778042/?report=abstract [Accessed January 25, 2021].

Love MI, Huber W, Anders S (2014) Moderated estimation of fold change and dispersion for RNA-seq data with DESeq2. Genome Biol 15:550 Available at: http://genomebiology.biomedcentral.com/articles/10.1186/s13059-014-0550-8 [Accessed February 14, 2020].

Luck K et al. (2019) A reference map of the human protein interactome. bioRxiv.

Marbach D, Lamparter D, Quon G, Kellis M, Kutalik Z, Bergmann S (2016) Tissue-specific regulatory circuits reveal variable modular perturbations across complex diseases. Nat Methods 13:366–370.

Mardinoglu A, Agren R, Kampf C, Asplund A, Nookaew I, Jacobson P, Walley AJ, Froguel P, Carlsson LM, Uhlen M, Nielsen J (2013) Integration of clinical data with a genome-scale metabolic model of the human adipocyte. Mol Syst Biol 9:649 Available at: https://onlinelibrary.wiley.com/doi/abs/10.1038/msb.2013.5 [Accessed January 17, 2020].

Mardinoglu A, Agren R, Kampf C, Asplund A, Uhlen M, Nielsen J (2014) Genome-scale metabolic modelling of hepatocytes reveals serine deficiency in patients with non-alcoholic fatty liver disease. Nat Commun 5:3083.

Mardinoglu A, Boren J, Smith U, Uhlen M, Nielsen J (2018) Systems biology in hepatology: approaches and applications. Nat Rev Gastroenterol Hepatol 15:365–377 Available at: http://www.nature.com/articles/s41575-018-0007-8 [Accessed November 21, 2019].

Martucci CP (1983) The role of 2-methoxyestrone in estrogen action. J Steroid Biochem 19:635–638.

McInnes L, Healy J, Saul N, Großberger L (2018) UMAP: Uniform Manifold Approximation and Projection. J Open Source Softw 3:861 Available at: http://joss.theoj.org/papers/10.21105/joss.00861 [Accessed January 20, 2021].

Meoni S, Macerollo A, Moro E (2020) Sex differences in movement disorders. Nat Rev Neurol 16:84–96 Available at: https://doi.org/10.1038/ [Accessed January 25, 2021].

Mostafavi S et al. (2018) A molecular network of the aging human brain provides insights into the pathology and cognitive decline of Alzheimer’s disease. Nat Neurosci 21:811–819.

Myers AJ et al. (2007) A survey of genetic human cortical gene expression. Nat Genet 39:1494–1499 Available at: https://www.nature.com/articles/ng.2007.16 [Accessed January 14, 2021].

Patil KR, Nielsen J (2005) Uncovering transcriptional regulation of metabolism by using metabolic network topology. Proc Natl Acad Sci U S A 102:2685–2689 Available at: www.pnas.orgcgidoi10.1073pnas.0406811102 [Accessed January 19, 2021].

Rajkumar AP, Bidkhori G, Shoaie S, Clarke E, Morrin H, Hye A, Williams G, Ballard C, Francis P, Aarsland D (2020) Postmortem Cortical Transcriptomics of Lewy Body Dementia Reveal Mitochondrial Dysfunction and Lack of Neuroinflammation. Am J Geriatr Psychiatry 28:75–86.

Rajsombath MM, Nam AY, Ericsson M, Nuber S (2019) Female Sex and Brain-Selective Estrogen Benefit α-Synuclein Tetramerization and the PD-like Motor Syndrome in 3K Transgenic Mice. J Neurosci 39:7628–7640 Available at: https://doi.org/10.1523/JNEUROSCI.0313-19.2019 [Accessed January 13, 2021].

Resnick SM et al. (2017) Testosterone treatment and cognitive function in older men with low testosterone and age-associated memory impairment. JAMA - J Am Med Assoc 317:717–727 Available at: /pmc/articles/PMC5433758/?report=abstract [Accessed January 21, 2021].

Riahi RR, Bush AE, Cohen PR (2016) Topical Retinoids: Therapeutic Mechanisms in the Treatment of Photodamaged Skin. Am J Clin Dermatol 17:265–276 Available at: https://pubmed.ncbi.nlm.nih.gov/26969582/ [Accessed January 21, 2021].

Ritchie ME, Phipson B, Wu D, Hu Y, Law CW, Shi W, Smyth GK (2015) Limma powers differential expression analyses for RNA-sequencing and microarray studies. Nucleic Acids Res 43:e47.

Sengoku R (2020) Aging and Alzheimer’s disease pathology. Neuropathology 40:22–29 Available at: https://onlinelibrary.wiley.com/doi/abs/10.1111/neup.12626 [Accessed January 25, 2021].

Shepardson NE, Shankar GM, Selkoe DJ (2011) Cholesterol level and statin use in Alzheimer disease: I. Review of epidemiological and preclinical studies. Arch Neurol 68:1239–1244 Available at: /pmc/articles/PMC3211071/?report=abstract [Accessed January 21, 2021].

Shudo K, Fukasawa H, Nakagomi M, Yamagata N (2009) Towards Retinoid Therapy for Alzheimers Disease. Curr Alzheimer Res 6:302–311 Available at: https://pubmed.ncbi.nlm.nih.gov/19519313/ [Accessed January 21, 2021].

Sinclair E, Trivedi DK, Sarkar D, Walton-Doyle C, Milne J, Kunath T, Rijs AM, de Bie RMA, Goodacre R, Silverdale M, Barran P (2021) Metabolomics of sebum reveals lipid dysregulation in Parkinson’s disease. Nat Commun 12:1–9 Available at: https://doi.org/10.1038/s41467-021-21669-4 [Accessed March 25, 2021].

Stampfer MJ (2006) Cardiovascular disease and Alzheimer’s disease: Common links. J Intern Med 260:211–223 Available at: http://doi.wiley.com/10.1111/j.1365-2796.2006.01687.x [Accessed March 24, 2020].

Supek F, Bošnjak M, Škunca N, Šmuc T (2011) Revigo summarizes and visualizes long lists of gene ontology terms Gibas C, ed. PLoS One 6:e21800 Available at: https://dx.plos.org/10.1371/journal.pone.0021800 [Accessed March 24, 2021].

Traag VA, Waltman L, van Eck NJ (2019) From Louvain to Leiden: guaranteeing well-connected communities. Sci Rep 9:1–12 Available at: https://doi.org/10.1038/s41598-019-41695-z [Accessed March 9, 2021].

Uhlén M et al. (2015) Tissue-based map of the human proteome. Science 347:1260419–1260419 Available at: http://www.sciencemag.org/cgi/doi/10.1126/science.1260419 [Accessed January 7, 2020].

Van Der Maaten L, Hinton G (2008) Visualizing data using t-SNE. J Mach Learn Res 9:2579–2625 Available at: http://jmlr.org/papers/v9/vandermaaten08a.html [Accessed January 20, 2021].

van Dijk D, Sharma R, Nainys J, Yim K, Kathail P, Carr AJ, Burdziak C, Moon KR, Chaffer CL, Pattabiraman D, Bierie B, Mazutis L, Wolf G, Krishnaswamy S, Pe’er D (2018) Recovering Gene Interactions from Single-Cell Data Using Data Diffusion. Cell 174:716–729.e27 Available at: https://doi.org/10.1016/j.cell.2018.05.061 [Accessed January 20, 2021].

Van Steijn L, Verbeek FJ, Spaink HP, Merks RMH (2019) Predicting Metabolism from Gene Expression in an Improved Whole-Genome Metabolic Network Model of Danio rerio. Zebrafish 16:348–362 Available at: https://pubmed.ncbi.nlm.nih.gov/31216234/ [Accessed January 19, 2021].

Väremo L, Nielsen J, Nookaew I (2013) Enriching the gene set analysis of genome-wide data by incorporating directionality of gene expression and combining statistical hypotheses and methods. Nucleic Acids Res 41:4378–4391 Available at: https://academic.oup.com/nar/article/41/8/4378/2408999 [Accessed January 18, 2021].

Wahjoepramono EJ, Asih PR, Aniwiyanti V, Taddei K, Dhaliwal SS, Fuller SJ, Foster J, Carruthers M, Verdile G, Sohrabi HR, Martins RN (2016) The Effects of Testosterone Supplementation on Cognitive Functioning in Older Men. CNS Neurol Disord - Drug Targets 15:337–343 Available at: /pmc/articles/PMC5078598/?report=abstract [Accessed January 21, 2021].

Wang H, Marcišauskas S, Sánchez BJ, Domenzain I, Hermansson D, Agren R, Nielsen J, Kerkhoven EJ (2018) RAVEN 2.0: A versatile toolbox for metabolic network reconstruction and a case study on Streptomyces coelicolor Ouzounis CA, ed. PLoS Comput Biol 14:e1006541 Available at: http://dx.plos.org/10.1371/journal.pcbi.1006541 [Accessed January 15, 2020].

Webster JA et al. (2009) Genetic Control of Human Brain Transcript Expression in Alzheimer Disease. Am J Hum Genet 84:445–458 Available at: http://www.cell.com/article/S0002929709001086/fulltext [Accessed January 14, 2021].

Wijemanne S, Jankovic J (2015) Dopa-responsive dystonia - Clinical and genetic heterogeneity. Nat Rev Neurol 11:414–424.

Wilkerson MD, Hayes DN (2010) ConsensusClusterPlus: A class discovery tool with confidence assessments and item tracking. Bioinformatics 26:1572–1573 Available at: https://pubmed.ncbi.nlm.nih.gov/20427518/ [Accessed January 18, 2021].

Yates AD et al. (2020) Ensembl 2020. Nucleic Acids Res 48:D682–D688 Available at: https://academic.oup.com/nar/article/48/D1/D682/5613682 [Accessed January 20, 2021].

Zhang Y, James M, Middleton FA, Davis RL (2005) Transcriptional analysis of multiple brain regions in Parkinson’s disease supports the involvement of specific protein processing, energy metabolism, and signaling pathways, and suggests novel disease mechanisms. Am J Med Genet - Neuropsychiatr Genet 137 B:5–16 Available at: https://pubmed.ncbi.nlm.nih.gov/15965975/ [Accessed July 9, 2020].

Zheng B et al. (2010) PGC-1α, a potential therapeutic target for early intervention in Parkinson’s disease. Sci Transl Med 2:52ra73 Available at: https://pubmed.ncbi.nlm.nih.gov/20926834/ [Accessed July 9, 2020].

